# Heme-Sensing Pathway Modulates Susceptibility of Poor Prognosis B-Lineage Acute Leukemia to BH3-Mimetics

**DOI:** 10.1101/2020.04.10.036319

**Authors:** Kaitlyn H. Smith, Amit Budhraja, John Lynch, Kathryn Roberts, John C. Panetta, Jon P. Connelly, Meghan E. Turnis, Shondra M. Pruett-Miller, John D. Schuetz, Charles G. Mullighan, Joseph T. Opferman

## Abstract

Anti-apoptotic *MCL1* is one of the most frequently amplified genes in human cancers and elevated expression confers resistance to many therapeutics including the BH3-mimetic agents ABT-199 and ABT-263. The anti-malarial, dihydroartemisinin (DHA) translationally represses MCL-1 and synergizes with BH3-mimetics. To explore how DHA represses MCL-1, a genome-wide CRISPR screen identified that loss of genes in the heme synthesis pathway renders mouse BCR-ABL^+^ B-ALL cells resistant to DHA-induced death. Mechanistically, DHA disrupts the interaction between heme and the eIF2α kinase heme regulated inhibitor (HRI) triggering the integrated stress response. Genetic ablation of *Eif2ak1*, which encodes HRI, blocks MCL-1 repression in response to DHA treatment and represses the synergistic killing of DHA and BH3-mimetics compared to wild-type leukemia. Furthermore, BTdCPU, a small-molecule activator of HRI, similarly triggers MCL-1 repression and synergizes with BH3-mimetics in mouse and human leukemia including both Ph^+^ and Ph-like B-ALL. Lastly, combinatorial treatment of leukemia bearing mice with both BTdCPU and a BH3-mimetic extended survival and repressed MCL-1 *in vivo*. These findings reveal for the first time that the HRI-dependent cellular heme-sensing pathway can modulate apoptosis in leukemic cells by repressing MCL-1 and increasing their responsiveness to BH3-mimetics. This signaling pathway could represent a generalizable mechanism for repressing MCL-1 expression in malignant cells and sensitizing them to available therapeutics.

## Introduction

While cure rates for children with B-precursor acute lymphoblastic leukemia (B-ALL) now exceed 90%, response rates in adults and children with poor prognosis B-ALL (e.g. *KMT2A* rearrangement, *BCR-ABL1* fusion, hypodiploid, etc.) are much poorer (1). Philadelphia chromosome positive (Ph^+^) B-ALL, which results from the fusion of the *BCR-ABL1* oncogene, encodes a constitutively active tyrosine kinase in B-ALL (2). Although treatment of these patients has improved with the use of tyrosine kinase inhibitors (TKI), many patients relapse and develop TKI resistance highlighting the need for improvements in treatment (3, 4). Another poor-prognosis subtype of ALL is Philadelphia chromosome like (Ph-like) ALL; Ph-like ALL has a gene expression profile similar to Ph^+^ B-ALL but is *BCR-ABL1* negative (5). Ph-like ALL is associated with high-risk clinical features and patients have a poorer response to chemotherapy than that of other subtypes (5, 6). Because Ph-like ALL involves a range of mutations leading to cytokine receptor and tyrosine kinase activations, precision medicine approaches have proven to be difficult and depend on the availability of targeted therapies (5). Therefore, new approaches are warranted to improve treatment for poor-prognosis B-ALL in adults and children.

Anti-apoptotic members of the BCL-2 family of proteins (e.g. BCL-2, BCL-XL, BFL-1, BCL-W, and MCL-1) are amplified in many types of cancers to prevent cell death (7, 8). *MCL1* is among the most frequently amplified genes in human cancer and many cancer cells, including B-ALL cells, depend on MCL-1 expression for survival (9, 10). As a result, so called BH3-mimetic drugs have been developed that inhibit anti-apoptotic members, allowing the release of pro-apoptotic members to trigger apoptosis (8). Navitoclax (ABT-263) which targets BCL-2/BCL-XL/BCL-W was the first-in-class BH3-mimetic, however, its clinical use has been hampered by dose limiting toxicities associated with on-target apoptosis induction in mature platelets (11). Currently, the only FDA approved BH3-mimetic is venetoclax (ABT-199), which specifically targets BCL-2 (12). Venetoclax has shown potency in a variety of malignancies, including chronic lymphocytic leukemia and acute myelogenous leukemia, especially when given in combination with standard chemotherapy (12–14). Venetoclax is well-tolerated in patients with low white blood cell counts as the most common side-effect (12). MCL-1 has been shown to be involved in resistance to ABT-199, as well as other chemotherapies, highlighting the need for MCL-1 targeting agents (15, 16). There has been notable progress in the clinical development of specific and potent MCL-1 inhibitors (e.g. S63845, AMG397, AMG176, and AZD5991); however, none are currently FDA approved, and the FDA recently placed a clinical hold on a phase I, dose escalation study of AMG397 after preliminary findings suggested some cardiac toxicity in patients (17–19). While the extent of the cardiac toxicity is still being evaluated, genetic deletion of *Mcl1* in adult mouse cardiomyocytes also triggered a rapid, fatal cardiomyopathy (20, 21).

Because direct pharmacological inhibition of MCL-1 may not be feasible due to on-target toxicities, it is important to investigate other mechanisms of targeting MCL-1. Our lab has previously shown that the anti-malarial drug dihydroartemisinin (DHA) can repress MCL-1 translation and synergize with the BH3-mimetic ABT-263 (22). Here, we show mechanistically that DHA induces the repression of MCL-1 through activation of a cellular heme-sensing pathway that results in eIF2α phosphorylation and subsequent attenuation of protein translation. The direct activation of this pathway by a small molecular tool compound can also repress MCL-1 and synergize with BH3-mimetics in both Ph^+^ and Ph-like leukemia. This observation reveals for the first time that this cellular heme-sensing pathway can be targeted to repress endogenous MCL-1 expression and lead to improved treatment of poor prognosis acute lymphoblastic leukemia.

## Materials and Methods

### Cells and Cell Culture

Mouse p185^+^ *Arf*-null B-ALL (hereafter referred to as BCR-ABL^+^ B-ALL) (10) and *PAX5-JAK2*, *RCSD1-AB1*, and *RCSD1-ABL2* fusion-expressing Ph-like Ba/F3 cells (23) were grown in RPMI with 10% fetal bovine serum, 55 µM 2-mercaptoethanol, 2 mM glutamine, penicillin, and streptomycin (Invitrogen). Navitoclax and Venetoclax were obtained from Selleckchem. DHA was obtained from AvaChem Scientific. BTdCPU was obtained from Millipore and synthesized by the Department of Chemical Biology and Therapeutics (St. Jude Children’s Research Hospital, SJCRH). The human Ph^+^ leukemia cell lines OP-1, TOM-1, BV-173, and SUP-B15 were cultured in RPMI with 20% fetal bovine serum, 55 µM 2-mercaptoethanol, 2 mM glutamine, penicillin, and streptomycin.

### Genome-wide CRISPR Screen

Cas9-expressing p185^+^ B-ALL cells were transduced with the Brie CRISPR KO library (Addgene #73633) at a multiplicity of infection of 0.5 with an sgRNA coverage of 400x (24). One day after transduction, puromycin was added (1 µg/mL) to select for infected cells. Cells were then cultured in either DMSO or 10 µM DHA containing media for 24 hours (h), followed by a 48h culture in drug free media. As previously described, genomic DNA was extracted from the surviving cells (Qiagen DNeasy Blood and Tissue kit) and amplified by PCR for Illumina sequencing of sgRNAs by Nextseq (24). MAGeCK was used to analyze the sequencing reads (25). Enrichr was used for pathway analysis of screen (26, 27).

### Validation CRISPR Screen

A validation CRISPR KO library was generated by the Center for Advanced Genome Editing (SJCRH) containing 5 guides (different from those in the Brie library) for each of the top 35 hits in positive selection (FDR <1%) and the top 13 hits in negative selection (FDR <16%) from the initial screen. Cas9-expressing BCR-ABL^+^ B-ALL cells were transduced with this validation library at a multiplicity of infection of 0.5 with a sgRNA coverage of >5000x. The drug treatment, sequencing and analysis were carried out in the same manner as the genome wide CRISPR screen.

### Knockout cell line creation

BCR-ABL^+^ B-ALL knockout cell lines were generated using CRISPR-Cas9 technology. Briefly, 1×10^6^ mouse BCR-ABL^+^ B-ALL cells were transiently co-transfected with precomplexed ribonuclear proteins (RNPs) consisting of 100 pmol of chemically modified sgRNA (Synthego) and 35 pmol of Cas9 protein (SJCRH Protein Production Core). Additionally, 200 ng of pMaxGFP was co-transfected via nucleofection (Lonza, 4D-Nucleofector™ X-unit) using solution P3 and program CM-137 in small cuvettes according to the manufacturer’s recommended protocol. Five days post nucleofection, cells were single-cell sorted by FACs to enrich for GFP^+^ (transfected) cells, clonally selected, and verified for the desired out-of-frame indel modifications via targeted deep sequencing on a Miseq Illumina sequencer. NGS analysis of clones was performed using CRIS.py (28). Two knockout clones for each gene were identified. BV-173 genetically modified cell pools were created by transiently co-transfecting 400,000 cells with precomplexed ribonuclear proteins (RNPs) consisting of 100 pmol of chemically modified sgRNA (Synthego) and 35 pmol of Cas9 protein (St. Jude Protein Production Core) via nucleofection (Lonza, 4D-Nucleofector™ X-unit) using solution P3 and program CA-137 in small cuvettes according to the manufacturers recommended protocol. Five days post nucleofection, a portion of cells were harvested and sequenced via targeted NGS and analyzed using CRIS.py as described above. NGS analysis indicated 89% total indels and 80% out-of-frame indels for the BV-173 edited cell pool. Sequences of sgRNA can be found in Sup. Table 1.

### Immunoblotting and Antibodies

Protein expression was assessed as previously described (29). Antibodies used were: anti-MCL-1 (Rockland Immunochemical), anti-human MCL-1, anti-PERK, anti-ATF6, anti-IRE1, anti-CHOP, anti-ATF4, anti-BCL-XL, anti-Phospho-eIF2α, anti-eIF2α (Cell Signaling), anti-HRI, and anti-Actin (Millipore) Anti-rabbit or anti-mouse horseradish peroxidase-conjugated secondary antibodies were from Jackson Immunochemical.

### Measuring Interaction between HRI and Heme

Absorption spectra were acquired using a Nanodrop 1000 spectrophotometer (Thermo Fisher) at room temperature. All dilutions were made in Enzyme Dilution Buffer (EDB) consisting of 50 mM Tris (pH 7.4), 150 mM NaCl, 0.1 mM EDTA, 25% glycerol. HRI (0.05 µg/ul) was obtained from ABM (Z500115). Heme (Frontier Scientific) and DHA were dissolved in DMSO to make an initial 10 mM stock, then further diluted in EDB to make 10X final stock immediately prior to addition. Enzyme was brought to 5 µM with heme and to the indicated concentration of DHA then incubated at room temperature for 30 minutes. Controls, without heme or DHA, had an equivalent volume of EDB added. All measurements were made after blanking with buffer (control) or with buffer containing the equivalent concentrations of heme and DHA as the measured sample. Absorbance readings between 220 and 750 nm were made for each sample, and Soret peak height for each independent experiment was determined by recording peak absorbance (420 nm) from 3 or more technical replicates per value.

### Heme measurements

Cellular heme measurements were made by reverse phase HPLC after pellets were extracted with acetone, acidified by the addition of 20% 1.6 N HCl, together with 10 pmol mesoporphyrin per sample added as an internal standard. Extracts were centrifuged at 21,100 × *g* for 10 min. after which pellets were discarded. From the supernatants, heme was separated from other porphyrins on a Shimadzu system (CBM-20A system controller, Shimadzu), using a mobile phase of acetonitrile in water containing 0.05% trichloroacetic acid at 1 ml/min on a reverse-phase C18 column (Sigma), applying a 30–66% linear acetonitrile gradient over 5 min followed by a 66– 90% linear gradient over 20 min. Heme was determined by measuring absorbance at 400 nm (Shimadzu, SPD-20AV). The concentration and identity of heme (t_R_=7.4 min.) was made by comparison with hemin (Frontier Scientific, H651-9) standards extracted analogously to samples. Calculations were made by normalizing the peak area for heme with internal standard peak values for samples then calculating from a linear curve made from hemin standards similarly normalized to the internal standard. Limit of detection was approximately 1 pmol with a linear range or detection to at least 1 nmol.

### Cell Death Experiments

Cells were seeded in 96-well plates and drugs (DHA, BTdCPU, ABT-199, ABT263 solubilized in DMSO or DMSO vehicle controls) were added at the indicated concentrations. Aminolevulinic acid (ALA), a precursor of heme synthesis, or succinylacetone (SA), an inhibitor of heme synthesis, (both from Sigma) were solubilized in water and added at indicated concentrations to alter cellular heme levels (30). Cell viability was determined by staining with Annexin-V-APC and propidium iodide (BD Biosciences) and measured by flow cytometry as previously described (22).

### Response Surface Modeling

Response surface modeling, implemented in Matlab version R2016a (Mathworks), was used to determine changes in the response of two drugs given in combination (31–33). A drug combination was considered either synergistic or antagonistic if the interaction term (α) describing the change in response relative to the additive model was either positive or negative, respectively. Two interaction terms (α) were considered different if their difference was statistically different from zero based on a two-tailed z-test.

### Patient-Derived Xenograft (PDX) Leukemia

Leukemia from adult patients with BCR-ABL1^+^ and *EBF1-PDGFRB* Ph-like ALL obtained from the Eastern Cooperative Oncology Group E2993 study (ClinicalTrials.gov identifier NCT00002514) and from the University Health Network (Toronto, CA) were transplanted into un-irradiated immunodeficient NOD.Cg-Prkdc^scid^Il2rg^tm1Wjl^/SzJ (NSG) mice (Jackson Laboratories) for 8-10 weeks prior to re-isolation (34–36). Mice were bred and utilized in accordance with SJCRH animal care and use committee (SJCRHACUC).

### Treatment of Leukemia in Recipient Mice

BCR-ABL^+^ B-ALL were injected (2×10^5^) into non-conditioned, 6-8-week-old, female C57BL/6 recipients (Jackson Laboratory). Five days after the transfer, recipients were treated with BTdCPU via intraperitoneal route and ABT-263 by oral gavage. Navitoclax was formulated in a mixture of 60% Phosal 50 PG, 30% PEG 400, and 10% EtOH and dosed at 100 mg/kg/day as previously described (37). 400 mg/kg/day of BTdCPU was administered in 30µL DMSO. Treatment was given daily for 14 days (days 5-18 after leukemia injection) during and after which the mice were monitored. Mice were bred and utilized in accordance with SJCRHACUC.

### Pathology and Immunohistochemistry

All tissues were fixed in formalin, embedded in paraffin, sectioned at 4 μm, mounted on positively charged glass slides (Superfrost Plus; Thermo Fisher Scientific), and dried at 60°C for 20 min before dewaxing and staining with hematoxylin and eosin (H&E) using standard methods. For immunohistochemical staining, the primary antibodies used in this study included anti-B220 (BD Biosciences) and anti-PAX5 (Abcam). Tissue sections underwent antigen retrieval in a prediluted Cell Conditioning Solution (CC1) (Ventana Medical Systems) for 32 min, and the OmniMap anti-Rabbit HRP kit (Ventana Medical Systems) and ChromoMap DAB (Ventana Medical Systems) were used for detection. All sections were examined by a pathologist blinded to the experimental group assignments.

## Results

### CRISPR screen identifies pathways required for DHA induced apoptosis

Previous efforts to mechanistically address how DHA triggered the repression of MCL-1 revealed a gene expression signature consistent with the induction of the endoplasmic reticulum (ER) stress pathway (22). To further interrogate this pathway, the three canonical branches of the cellular ER stress pathway were genetically ablated in mouse p185^+^ *Arf*-null B-ALL (hereafter referred to as BCR-ABL^+^ B-ALL) using CRISPR/Cas9 targeting (Sup. Fig. 1A). Despite loss of the genes encoding the IRE1, ATF6, or PERK branches of the ER stress pathway, DHA treatment still triggered MCL-1 repression in mouse BCR-ABL^+^ B-ALL, indicating that none of the canonical ER stress pathway signaling arms is singularly responsible for repressing MCL-1 expression in response to DHA treatment (Sup. Fig. 1A).

To elucidate which cellular signaling pathways are activated by DHA to induce cell death in mouse BCR-ABL^+^ B-ALL cells, an unbiased genetic screen was conducted in which Cas9-expressing mouse BCR-ABL^+^ B-ALL cells were stably transduced with the Brie knockout library (78,637 sgRNAs targeting 19,674 genes) (24). The cells were treated with 10 µM DHA for 24 hours (h) and cultured in drug free media for an additional 48h to allow outgrowth of resistant cells (Fig. 1A). After DHA treatment, BCR-ABL^+^ B-ALL cell viability was approximately 40% in contrast to vehicle control treated cells that maintained >95% cell viability (Fig. 1B). After treatment, genomic DNA was isolated from the viable BCR-ABL^+^ B-ALL cells, amplified and subjected to Illumina sequencing to determine sgRNAs enrichment in the surviving cells. MAGeCK analysis of sequencing data identified 35 genes whose targeting sgRNAs were enriched in DHA-resistant cells at a false discovery rate of <1% (25), indicating that loss of expression of these genes generated resistance to 10 µM DHA (Fig. 1C and Table 1). Enrichr gene set enrichment analysis was performed (26, 27) using these 35 genes, to identify genetic pathways implicated in DHA resistance. Multiple pathways related to heme synthesis/metabolism and apoptosis were identified (Fig. 1D and Table 1); these data implicate that either the heme synthesis or intrinsic apoptotic pathway is required for induction of apoptosis in BCR-ABL^+^ B-ALL cells in response to 10 µM DHA.

**Figure 1.**
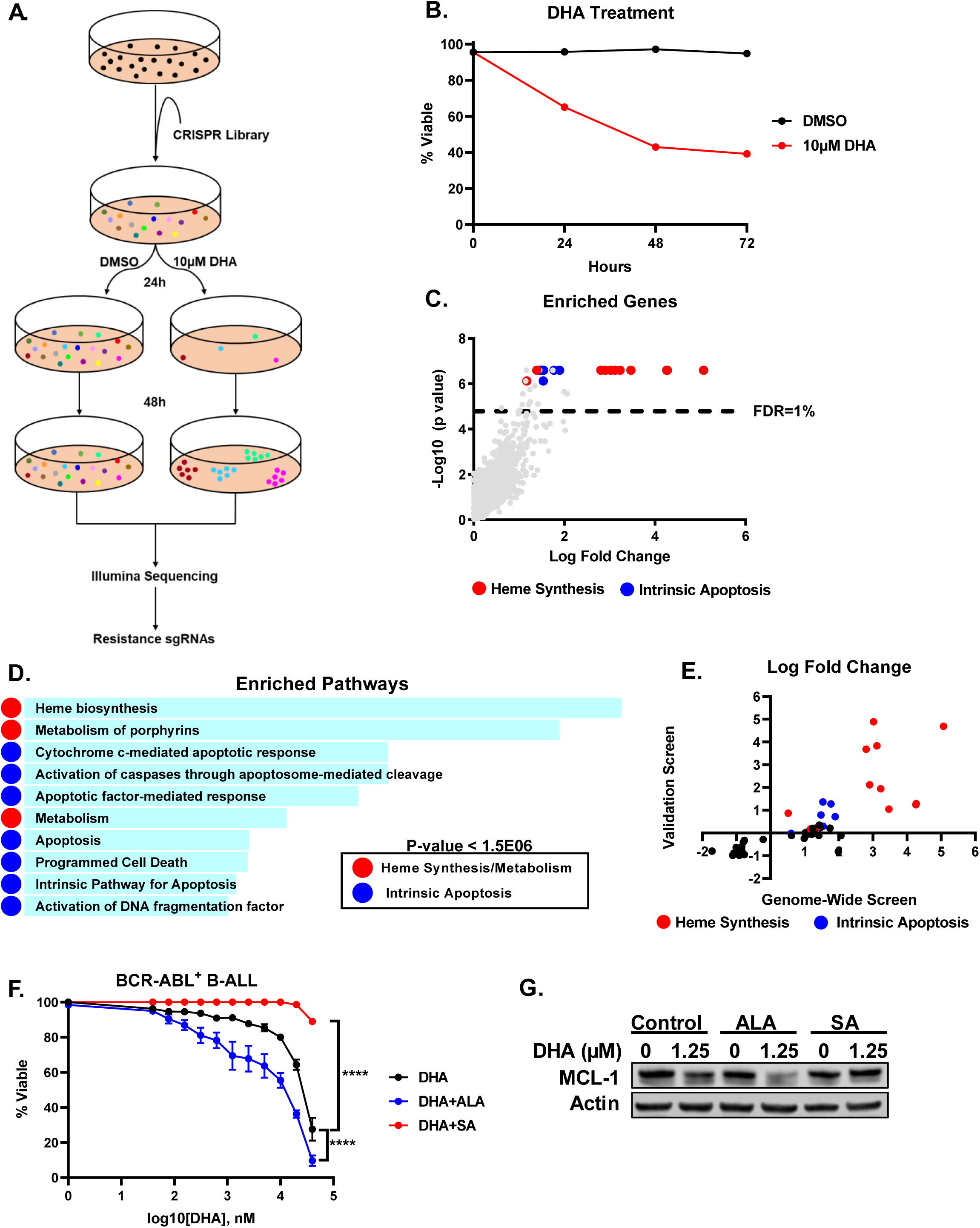
CRISPR screen identifies the requirement of the heme synthesis pathway for DHA induced apoptosis. (**A**) BCR-ABL^+^ B-ALL cells containing the Brie CRISPR knockout library were treated with DHA or DMSO for 24h and then cultured for an additional 48h in drug free media. Cells that survived the drug treatment were harvested and sequenced to determine genes involved in resistance. (**B**) Percentage of viable cells treated with either 10 µM DHA or DMSO was measured by trypan blue staining. Cells were under treatment conditions for 24h and then cultured in drug free media for an additional 48h. (**C**) MAGeCK was used to determine sgRNAs enriched for in the DHA treated sample. (**D**) Enrichr was used to determine pathways involved in resistance to DHA. **(E)** BCR-ABL^+^ B-ALL cells containing the validation screen CRISPR library were treated with DHA for 24h and cultured for an additional 48h in drug free media. Surviving cells were harvested and the DNA sequenced. MAGeCK was used to determine sgRNAs enriched for in the DHA treated sample. The enriched genes from both the validation screen and original genome-wide screen were then compared. (**F and G**) Cells were treated with a combination of DHA and 62.5 µM of Aminolevulinic acid (ALA) or succinylacetone (SA) for 24h. (**F**) Viability was measured using Annexin-V and propidium iodide staining. Data are the average of three experiments and error bars are SEM. Two-way ANOVA with Bonferroni multiple comparison indicates significance P<0.0001**** between the DHA alone and DHA+ALA or DHA+SA treatments. (**G**) Protein expression was determined by immunoblotting lysates from (F) with indicated antibodies.

**Table 1.**
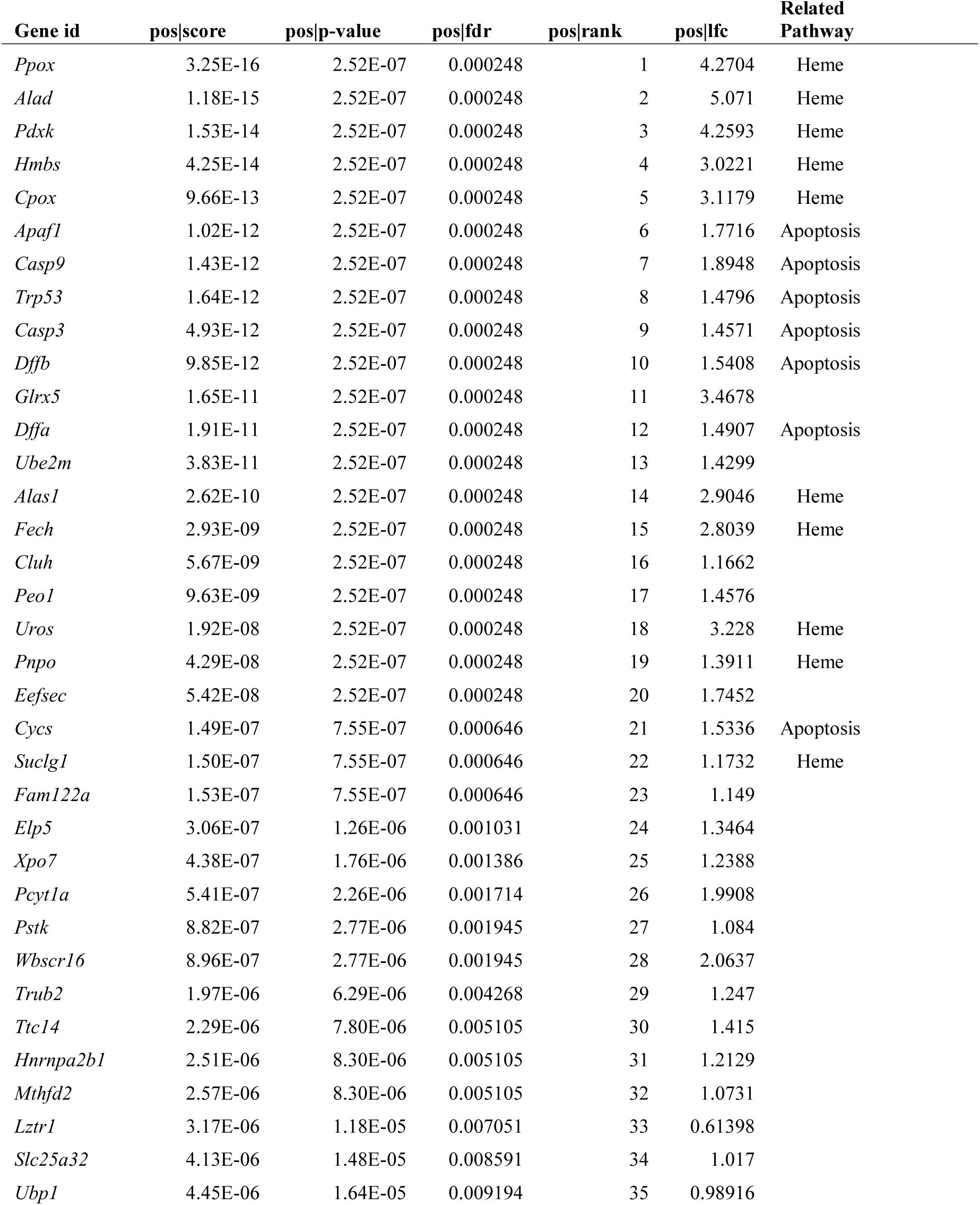
Top 35 hits from genome wide CRISPR screen, determined by MAGeCK. Pos|Score: The RRA value of the gene in positive selection. Pos | p-value: raw p-value of this gene in positive selection. Pos | fdr: false discovery rate of this gene in positive selection. Pos | rank: ranking of each gene in positive selection Pos | lfc: log fold change of the gene in positive selection. Related Pathway: category of gene as determined by Enrichr/GSEA.

To validate the hits identified from the primary genome-wide screen a secondary targeted screen was performed using a validation library composed of five, new sgRNA guides per gene for the top 35 hits in positive selection (FDR <1%) and the top 13 hits in negative selection (FDR <16%). The validation screen was carried out in the same way as the original genome-wide screen. After MAGeCK analysis, 8 of the 10 genes involved in heme synthesis and 4 of the 7 apoptosis-related genes identified in the genome-wide screen were found to be top hits in the validation screen (FDR<12%) (Fig. 1E). These data strongly suggest that DHA requires the heme synthesis pathway to kill BCR-ABL^+^ B-ALL cells.

Since sgRNAs targeting the heme synthesis pathway were associated with resistance of BCR-ABL^+^ B-ALL cells to DHA treatment, we hypothesized that repressing heme levels pharmacologically should also render BCR-ABL^+^ B-ALL cells resistant to DHA treatment. To test this hypothesis, we cultured BCR-ABL^+^ B-ALL cells in succinylacetone (SA), an inhibitor of heme synthesis (30), resulting in decreased cellular heme levels and rendering the cells more resistant to death induced by culture with DHA (Sup. Fig. 1B and Fig. 1F). Conversely, BCR-ABL^+^ B-ALL cells were cultured in aminolevulinic acid (ALA), a precursor of heme synthesis (30), which resulted in increased cellular heme levels and made the cells more sensitive to DHA induced apoptosis (Sup. Fig. 1B and Fig. 1F). Since DHA acts to repress MCL-1 translation in leukemic cells (22), we tested whether pharmacological manipulation of cellular heme levels could affect MCL-1 repression in response to DHA treatment. The combination of ALA and DHA caused even greater MCL-1 repression than that observed with DHA alone (Fig. 1G). In contrast, the inhibition of heme synthesis by SA substantially blunted MCL-1 repression triggered by DHA (Fig. 1G). Taken together, these data indicate that cellular heme is needed both for DHA to induce apoptosis and to repress MCL-1 expression.

### DHA induces an HRI mediated eIF2α phosphorylation

The observation that heme is required for DHA induced apoptosis and repression of MCL-1 prompted us to further examine how the heme pathway could trigger the translational repression of MCL-1. The phosphorylation of eIF2α inhibits cap-dependent translation by triggering the integrated stress response (38). Indeed, when BCR-ABL^+^ B-ALL cells were treated with DHA, eIF2α was phosphorylated and the protein expression of both ATF4 and the CHOP transcription factor were induced (Fig. 2A). Concomitantly, MCL-1 expression was repressed, but the expression of other anti-apoptotic BCL-2 family members, such as BCL-XL, was unaffected (Fig. 2A). There are four known eIF2α kinases, GCN2, PKR, PERK, and the heme regulated inhibitor (HRI) (39). PERK-deficient cells still repressed MCL-1 in response to DHA treatment (Sup. Fig. 1A), thus excluding PERK and the classical ER stress pathway as being responsible for eIF2α phosphorylation triggered by DHA.

**Figure 2.**
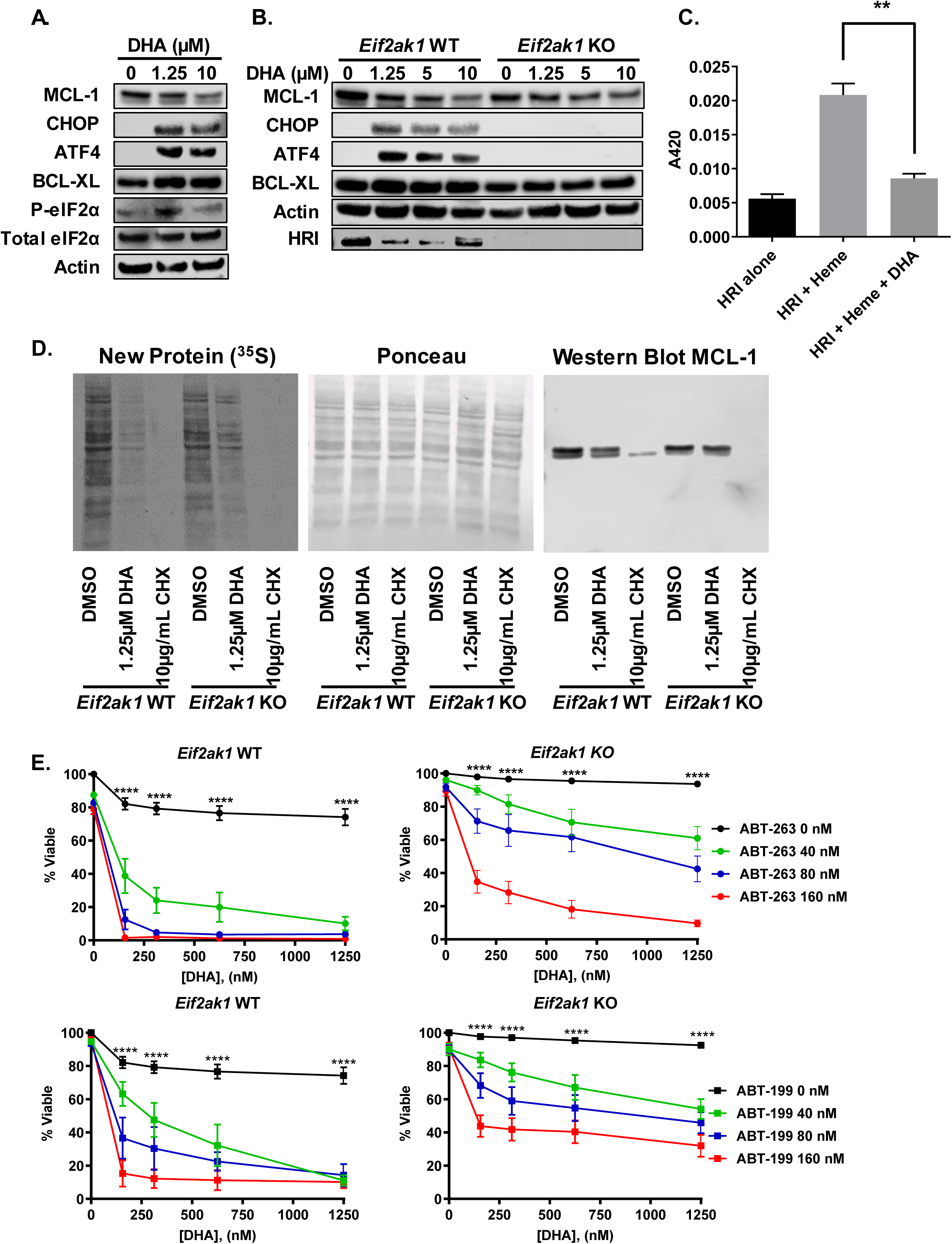
HRI is required for DHA induced MCL-1 repression and synergism with BH3-mimetics. (**A**) BCR-ABL^+^ B-ALL cells were treated with DHA for 16h and protein expression was determined by immunoblotting with indicated antibodies. **(B)** Wild-type (WT) or *Eif2ak1*-KO (lacking HRI) BCR-ABL^+^ B-ALL cells were treated with DHA for 16h and protein expression was determined by immunoblotting with indicated antibodies. **(C)** Purified HRI protein was incubated with 20 µM heme, 20 µM DHA, or a combination of the two at room temperature for 30 min. Absorbance was then measured at 420 nm to detect the presence of a Soret peak which is indicative of protein binding. Data are the average of three experiments and error bars are SEM. Unpaired t-test indicates significance between HRI+Heme vs. HRI+Heme+DHA p<0.01**. **(D)** Wild-type or *Eif2ak1*-KO cells were treated with the indicated drugs for 9h and then pulse-labeled with ^35^S-methionine-cysteine for 1h. Autoradiography was used to determine new protein synthesis. Cycloheximide (CHX) served as a positive control for translation inhibition. Total protein was determined by Ponceau staining. MCL-1 expression was determined by immunoblotting. **(E)** WT or *Eif2ak1*-KO BCR-ABL^+^ B-ALL cells were treated with ABT-263 or ABT-199 (0, 40, 80, 160 nM) alone or in combination with the indicated concentrations of DHA for 24h. Viable cells were measured using Annexin-V and propidium iodide staining. Data are the average of three experiments and error bars are SEM. Two-way ANOVA with Bonferroni multiple comparison indicates significance P<0.0001**** between the DHA alone (0 nM ABT-263 or ABT-199) and 160 nM ABT-263 or ABT-199 at indicated doses of DHA. The combination of DHA+ABT-263 showed a statistically less synergistic response in *Eif2ak1*-KO (α=3.15, p=5.67e-06) as compared to wild-type (α=5.41, p=7.5e-08) BCR-ABL^+^ B-ALL cells (p<10^-5^). The combination of DHA+ABT-199 showed a significantly less synergistic response in *Eif2ak1*-KO (α=5.4, p=1.16e-83) as compared to wild-type (α=7.96, p=2.7e-32) BCR-ABL B-ALL cells (p<0.057).

The eIF2α kinase HRI phosphorylates eIF2α in erythrocytes to prevent hemoglobin translation when cellular heme levels are limiting (40). To assess whether HRI is responsible for MCL-1 repression in BCR-ABL^+^ B-ALL cells treated with DHA, we generated *Eif2ak1*-deficient (*Eif2ak1* encodes HRI, hereafter referred to as HRI-deficient) BCR-ABL^+^ B-ALL cells using CRISPR/Cas9. In contrast to wild-type BCR-ABL^+^ B-ALL cells, HRI-deficient BCR-ABL^+^ B-ALL cells, failed to induce the expression of either ATF4 or CHOP when treated with DHA (Fig. 2B). Notably, while DHA treatment in HRI expressing BCR-ABL^+^ B-ALL cells triggered the repression of MCL-1 expression, the BCR-ABL^+^ B-ALL cells lacking HRI failed to repress MCL-1 expression (Fig. 2B and D). These data indicate that HRI is responsible for the DHA induced eIF2α phosphorylation and activation of the down-stream cellular stress response, which represses MCL-1 expression.

In erythrocytes, HRI directly binds to heme, which represses kinase activity; however, when heme levels drop, HRI becomes active and phosphorylates eIF2α (41, 42). To address how DHA triggers HRI activation, we assessed the HRI-heme interaction using spectroscopy on purified, recombinant proteins. When HRI and heme were incubated together, a Soret peak was detected indicating binding of the two proteins. However, when DHA was incubated with heme and HRI, the Soret peak was lost indicating that DHA addition disrupts the interaction between heme and HRI (Fig. 2C). These data indicate that DHA can disrupt the HRI-heme interaction in a cell-free system.

### HRI is required for DHA induced MCL-1 translational repression and the synergistic response of DHA and BH3-mimetics

HRI-mediated phosphorylation of eIF2a represses cap-dependent translation and blocks a wide array of protein synthesis (38). To confirm whether active HRI is upstream of the MCL-1 translational repression we used metabolic labeling to assess whether loss of HRI attenuates translational inhibition triggered by DHA treatment. HRI wild-type or deficient BCR-ABL^+^ B-ALL cells were treated with DHA for 9h and then pulse-labeled with 100 µCi/ml of ^35^S-labeled methionine and cysteine for 1h. Autoradiography of whole cell lysates from treated cells demonstrated that global translation was repressed in DHA treated control cells; however, the repression of translation was substantially blunted in HRI-deficient cells in response to DHA treatment (Fig. 2D). As a positive control, culture of both cell types with cycloheximide (CHX) repressed translation in an HRI-independent manner. Western blotting of the treated cells also revealed that MCL-1 was repressed in the control cells, but not the HRI-deficient BCR-ABL^+^ B-ALL cells (Fig. 2D). These data indicate that DHA treatment induces an HRI-dependent inhibition of translation that results in MCL-1 repression.

Our previous studies demonstrated that DHA-induced repression of MCL-1 renders BCR-ABL^+^ B-ALL cells susceptible to synergistic cell death induced by the BH3-mimetic drug ABT-263 (22). Since HRI is responsible for mediating the repression of MCL-1 in response to DHA treatment, we assessed whether HRI-deficient BCR-ABL^+^ B-ALL cells would exhibit less synergistic cell death in response to combination treatment of DHA with BH3-mimetic drugs. In control BCR-ABL^+^ B-ALL cells DHA synergized with both BH3-mimetic drugs ABT-199 and ABT-263 as evidenced by response surface modeling (Fig. 2E and Sup. Fig. 2). However, the loss of HRI from BCR-ABL^+^ B-ALL cells substantially blunted the synergistic killing between DHA and both BH3-mimetics, indicating that loss of HRI attenuated the synergism between DHA and BH3-mimetics because MCL-1 was not repressed (Fig. 2E and Sup. Fig. 2).

### Direct activation of HRI represses MCL-1 and synergizes with BH3-mimetics

DHA repressed MCL-1 in an HRI-dependent manner, so we asked whether a small molecule activator of HRI, BTdCPU (43), could also induce the repression of MCL-1. Similar to DHA treatment, when BCR-ABL^+^ B-ALL cells were treated with BTdCPU MCL-1 expression was repressed whereas BCL-XL expression was unaffected (Fig. 3A). Consistently, much less MCL-1 repression was induced by BTdCPU treatment in the HRI-deficient leukemic cells indicating that BTdCPU induced an HRI-dependent repression of MCL-1 (Fig. 3A). Additionally, BTdCPU triggered the integrated stress response and induced the downstream targets CHOP and ATF4 in wild-type, but not HRI-deficient BCR-ABL^+^ B-ALL cells (Fig. 3A). These data indicate that BTdCPU can activate HRI and induce MCL-1 repression.

**Figure 3.**
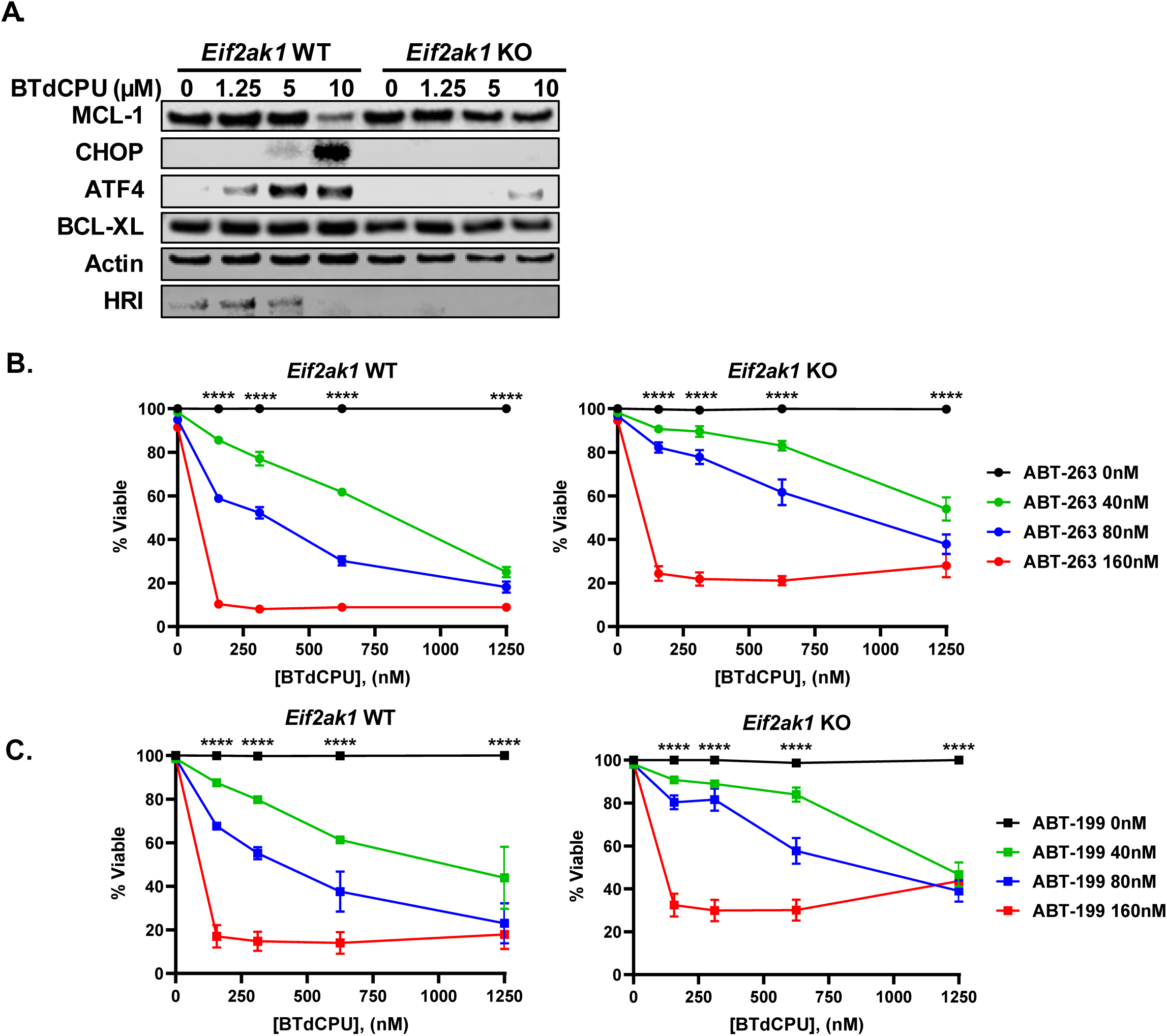
Direct activation of HRI represses MCL-1 and synergizes with BH3-mimetics. **(A)** Wild-type (WT) or *Eif2ak1*-KO (HRI-deficient) BCR-ABL^+^ B-ALL cells were treated with BTdCPU for 16h and protein expression was determined by immunoblotting with indicated antibodies. (**B and C**) WT or *Eif2ak1*-KO BCR-ABL^+^ B-ALL cells were treated with **(B)** ABT-263 or **(C)** ABT-199 (0, 40, 80, 160 nM) alone or in combination with the indicated concentrations of BTdCPU for 24h. Viable cells were measured using Annexin-V and propidium iodide staining. Data are the average of three experiments and error bars are SEM. Two-way ANOVA with Bonferroni multiple comparison indicates significance P<0.0001**** between the BTdCPU alone (0 nM ABT-263 or ABT-199) and 160 nM ABT-263 or ABT-199 at indicated doses of BTdCPU. The combination of BTdCPU+ABT-263 showed a statistically less synergistic response in *Eif2ak1*-KO (α=0.435, p=1.16e-12) as compared to wild-type (α=0.66, p=2.826e-13) BCR-ABL^+^ B-ALL cells (p<0.05). The combination of BTdCPU+ABT-199 showed a statistically less synergistic response in *Eif2ak1*-KO (α=1.66, p=3.79e-18) as compared to wild-type (α=2.27, p=1.75e-26) BCR-ABL B-ALL cells (p<0.05).

Since BTdCPU can induce MCL-1 repression in an HRI-dependent manner, we assessed whether it can synergize with BH3-mimetics in killing BCR-ABL^+^ B-ALL cells. When BTdCPU was combined with either ABT-199 or ABT-263 it induced a synergistic cell death response in wild-type BCR-ABL^+^ B-ALL cells (Fig. 3B-C and Sup. Fig. 3). However, in HRI-deficient BCR-ABL^+^ B-ALL cells the synergistic combination of BTdCPU with both BH3-mimetics was significantly attenuated (Fig. 3B-C and Sup. Fig. 3). These data indicate that the activation of HRI by BTdCPU can induce synergistic leukemia cell killing in our murine cell model of BCR-ABL^+^ B-ALL.

### HRI activation synergizes with BH3-mimetics in ALL cell lines

To assess whether activation of HRI can synergize with BH3-mimetic agents in human Ph^+^ leukemia cells, BV-173 cells, were cultured with either DHA or BTdCPU. Similar to our observations in mouse BCR-ABL^+^ B-ALL cells, treatment of human leukemia cells with DHA or BTdCPU induced eIF2α phosphorylation, expression of CHOP and ATF4, and repression of MCL-1 without altering BCL-XL expression (Fig. 4A). In murine BCR-ABL^+^ B-ALL cells, the repression of MCL-1 triggered by either DHA or BTdCPU required the HRI-mediated activation of the integrated stress response; therefore, we sought to determine whether human Ph^+^ acute leukemia cells were similarly HRI-dependent. To this aim, *EIF2AK1* (encoding HRI) was ablated in BV-173 Ph^+^ leukemia cell lines by CRISPR/Cas9 and the cells were subjected to treatment with DHA, BTdCPU, or thapsigargin (an inducer of the classical ER stress pathway) (44). As in mouse BCR-ABL^+^ B-ALL cells, loss of HRI expression prevented the repression of MCL-1 expression in cells cultured with DHA or BTdCPU and blocked the repression of ATF4 and CHOP (Fig. 4B). In contrast, thapsigargin treatment still repressed MCL-1 and induced ATF4 and CHOP in BV-173 cells lacking HRI, confirming that the ER stress pathway does not depend on HRI (Fig. 4B). These data indicate that activation of HRI-mediated eIF2α phosphorylation pathway, by either DHA or BTdCPU treatment, can cause MCL-1 repression and therefore should be able to synergize with BH3-mimetics in human Ph^+^ leukemia cells. Consistent with these findings, when BV-173 cells were cultured with either DHA or BTdCPU and combined with BH3-mimetics (ABT-199 or ABT-263), the combined treatments were significantly synergistic with both BH3-mimetics, but in BV-173 cells lacking HRI expression the synergism was significantly reduced (Fig. 4C and Sup. Fig. 4A-B). Similar synergism between either DHA or BTdCPU and BH3-mimetics were obtained in human TOM1, OP-1, and SUP-B15 Ph^+^ leukemia cell lines treated in culture (Sup. Fig. 4C-E and Sup. Table 2).

**Figure 4.**
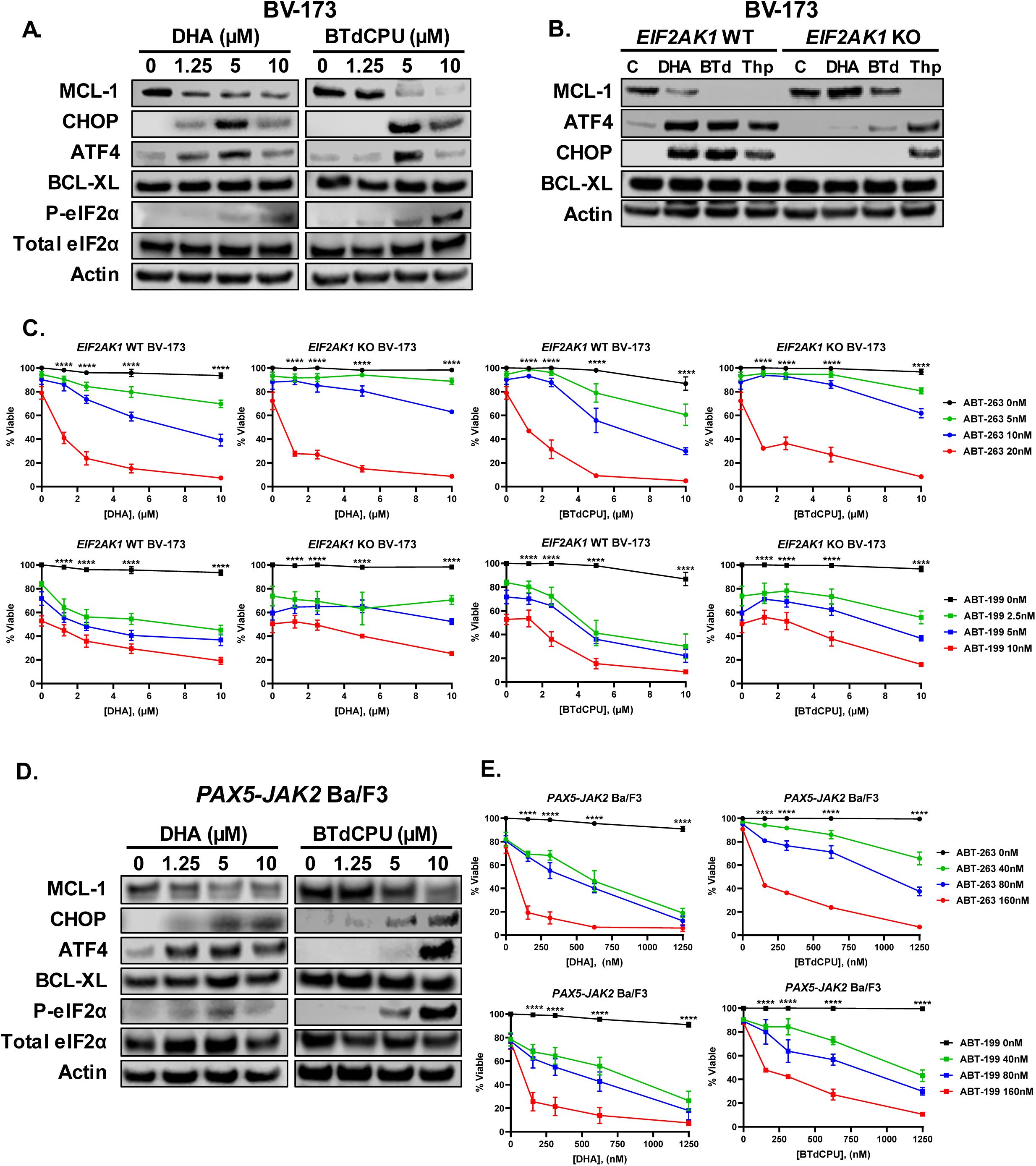
HRI activation represses MCL-1 and synergizes with BH3-mimetics in Ph^+^ and Ph-like ALL cell lines. (**A**) Human Ph^+^ BV-173 cells were treated with the indicated concentrations of DHA or BTdCPU for 24h and protein expression was determined by western blotting. (**B**) Wild-type (WT) or a knockdown pool of *EIF2AK1*-KO (lacking HRI) BV-173 cells were treated with DMSO (Control, C), DHA (5 µM), BTdCPU (BTd, 5 µM), or Thapsigargin (Thp, 100 nM) for 24h and protein expression was determined by western blotting. **(C)** WT or *EIF2AK1*-KO BV-173 cells were treated with ABT-263 (0, 5, 10, 20 nM) or ABT-199 (0, 2.5, 5, 10 nM) alone or in combination with the indicated concentrations of DHA or BTdCPU for 24h. Viable cells were measured using Annexin-V and propidium iodide staining. Data are the average of 3 experiments and error bars are SEM. Two-way ANOVA with Bonferroni multiple comparison indicates significance P<0.0001**** between DHA or BTdCPU alone (0 nM ABT-263 or ABT-199) and 20 nM ABT-263 or 10 ABT-199 at indicated doses of DHA or BTdCPU. The combination of DHA+ABT-263 showed significantly less synergistic response in BV-173 *EIF2AK1*-KO (α=23, p=1.05e-03) as compared to BV-173 wild-type (α=94.2, p=1.66e-08) cells (p<0.003). The combination of DHA+ABT-199 showed a less synergistic response in BV-173 *EIF2AK1*-KO (α=1.58, p=2.54e-01) as compared to BV-173 wild-type (α=69.1, p=4.49e-08) cells (p<0.003). The combination of BTdCPU+ABT-199 showed a less synergistic response in *EIF2AK1*-KO (α=3.33, p=3.05e-02) as compared to wild-type (α=12.3, p=5.53e-06) cells (p<0.003). (**D**) *PAX5-JAK2* Ba/F3 cells were treated with the indicated doses of DHA or BTdCPU for 9h and protein expression was determined by immunoblotting with indicated antibodies. (**E**) *PAX5-JAK2* Ba/F3 cells were treated with the indicated drugs for 24h. Viable cells were measured using Annexin-V and propidium iodide staining. Data are the average of three experiments and error bars are SEM. Two-way ANOVA with Bonferroni multiple comparison indicates significance P<0.0001**** between DHA or BTdCPU alone (0 nM ABT-263 or ABT-199) and 160 nM ABT-263 or ABT-199 at indicated doses of DHA or BTdCPU. The combination of DHA+ABT-263 or ABT-199 showed synergistic response in Ba/F3 cells (α=220, p=1.89e-33) and (α=202, p=2.48e-34) respectively (upper panel). The combination of BTdCPU+ABT-263 or ABT-199 showed synergistic response in Ba/F3 cells (α=98.2, p=1.14e-56) and (α=129, p=6.59e-52) respectively (lower panel).

We next wanted to extend our findings to other poor prognosis subtypes of ALL. To achieve this aim, we tested DHA and BTdCPU in a mouse cell line expressing a fusion oncoprotein driver of Ph-like ALL. Mouse Ba/F3 cells stably expressing a human *PAX5-JAK2* fusion were treated with DHA or BTdCPU to determine if MCL-1 was repressed (23). Consistent with Ph^+^ leukemia cell lines, we found that MCL-1 expression was repressed in response to DHA or BTdCPU in *PAX5-JAK2* Ba/F3 cells (Fig. 4D). Additionally, eIF2α phosphorylation was induced, ATF4 and CHOP were expressed, and BCL-XL protein was unaffected by treatment with either DHA or BTdCPU. Since MCL-1 was repressed in the *PAX5-JAK2* Ba/F3 cells, we next investigated whether there was a synergistic effect when DHA or BTdCPU was combined with BH3-mimetics. As expected, synergy was observed in the *PAX5-JAK2* Ba/F3 cells when DHA or BTdCPU was combined with BH3-mimetics (Fig. 4E and Sup. Fig. 4F-G). In other Ba/F3 cells expressing the *RCSD1-ABL1* or *RCSD1-ABL2* Ph-like fusions (23); both DHA and BTdCPU also synergized with BH3-mimetics (Sup. Fig. 4H-I and Sup. Table 2).

### HRI activation synergizes with BH3-mimetics in primary patient-derived xenografts of human B-ALL

To confirm whether DHA or BTdCPU can lead to synergistic responses when combined with ABT-199 or ABT-263 BH3-mimetics in primary patient leukemia, we took advantage of primary patient derived xenografts (PDXs) established from patients with Ph^+^ or *EBF1-PDGFRB* expressing Ph-like B-ALL. PDX cells isolated from recipient mice were cultured overnight with either DHA or BTdCPU and either ABT-199 or ABT-263 BH3-mimetics. Similar to Ph^+^ human cell lines, the combination of either BH3-mimetic agent synergized with both DHA and BTdCPU in Ph^+^ B-ALL patient derived xenograft samples (Fig. 5A and Sup. Fig. 5A). Consistently, *EBF1-PDGFRB* Ph-like PDX cells also responded synergistically to the combination of DHA or BTdCPU with either BH3-mimetic (Fig. 5B and Sup. Fig. 5B). These data indicate that primary patient leukemic cells can respond to either DHA or BTdCPU-induced synergism with BH3-mimetics agents in culture.

**Figure 5.**
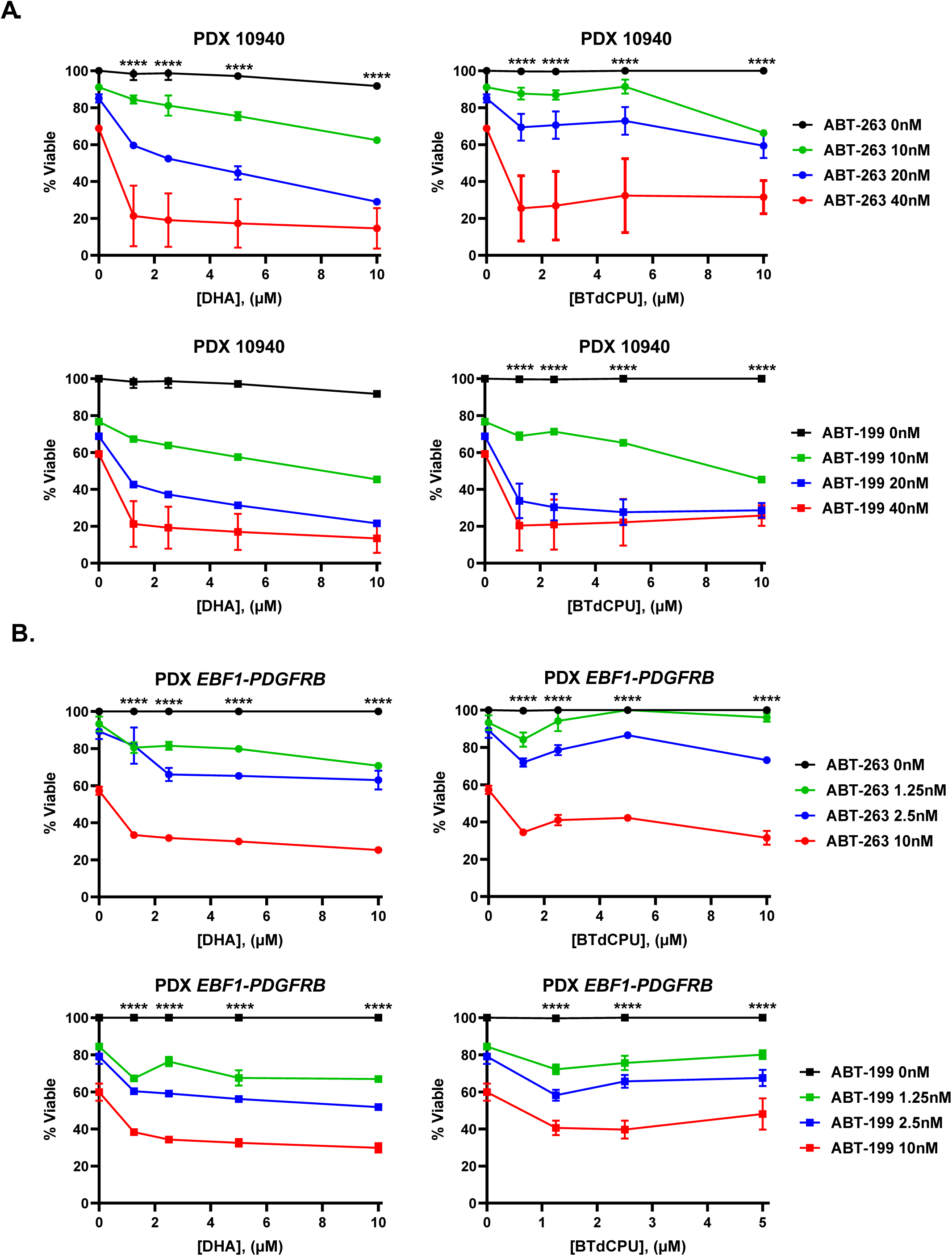
HRI activation synergizes with BH3-mimetics in Ph^+^ and Ph-like PDX cells. (**A**) Ph^+^ 10940 or (**B**) *EBF1-PDGFRB* Ph-like patient derived xenograft (PDX) cells were treated with the indicated concentrations of ABT-263 or ABT-199 alone or in combination with DHA or BTdCPU for 16h. Viable cells were measured using Annexin-V and propidium iodide staining. Data are the average of four experiments and error bars are SEM. Two-way ANOVA with Bonferroni multiple comparison indicates significance P<0.0001**** between DHA or BTdCPU alone (0 nM ABT-263 or ABT-199) and (40 nM ABT-263 or ABT-199) **(for A)** or (10 nM ABT-263 or ABT-199) at indicated doses of DHA or BTdCPU **(for B). (A)** The combination of DHA+ABT-263 or ABT-199 showed synergistic response in Ph^+^ PDX cells (α=1.3, p=2.11e-08) and (α=1.28, p=1.15e-07) respectively. The combination of BTdCPU+ABT-263 or ABT-199 showed synergistic response in Ph^+^ PDX cells (α=0.672, p=9.34e-03) and (α=0.98, p=2.45e-05) respectively. **(B)** The combination of DHA+ABT-263 or ABT-199 showed synergistic response in Ph-like cells (α=0.739, p=6.89e-10) and (α=0.629, p=1.66e-12) respectively. The combination of BTdCPU+ABT-263 orABT-199 showed synergistic response in Ph-like cells (α=0.432, p=1.37e-04) and (α=0.18, p=5.69e-04) respectively.

### HRI activation represses MCL-1 and synergizes with BH3-mimetics *in vivo*

Since combining HRI activation by BTdCPU synergized with BH3-mimetics in cultured mouse and human leukemia, we hypothesized that combining BTdCPU with ABT-263 could prolong survival of mice bearing BCR-ABL^+^ B-ALL. To this aim, C57/BL6 mice were injected with BCR-ABL^+^ B-ALL cells followed by treatment on day 5 after transplant with either vehicle, BTdCPU (400 mg/kg), ABT-263 (100 mg/kg), or a combination of BTdCPU and ABT-263. Treatments continued daily for 14 days during which mice were monitored. Recipient mice receiving vehicle or either BTdCPU or ABT-263 alone exhibited rapidly progressing leukemia and all mice from these cohorts required euthanasia within 13 days after leukemia injection due to progressive leukemia irrespective of treatment regimen. In contrast, recipient mice treated with the combination of BTdCPU and ABT-263 for 14 days exhibited significantly prolonged survival (Fig. 6A). Complete blood counts from peripheral blood on day 9 revealed that mice receiving vehicle or BTdCPU alone had higher white blood cell counts (WBCs) whereas those mice receiving the combination treatment had statistically lower WBCs than vehicle treated recipients (Fig. 6B). These data indicated that the combination of BTdCPU and ABT-263 reduced the leukemia burden in recipient mice.

**Figure 6.**
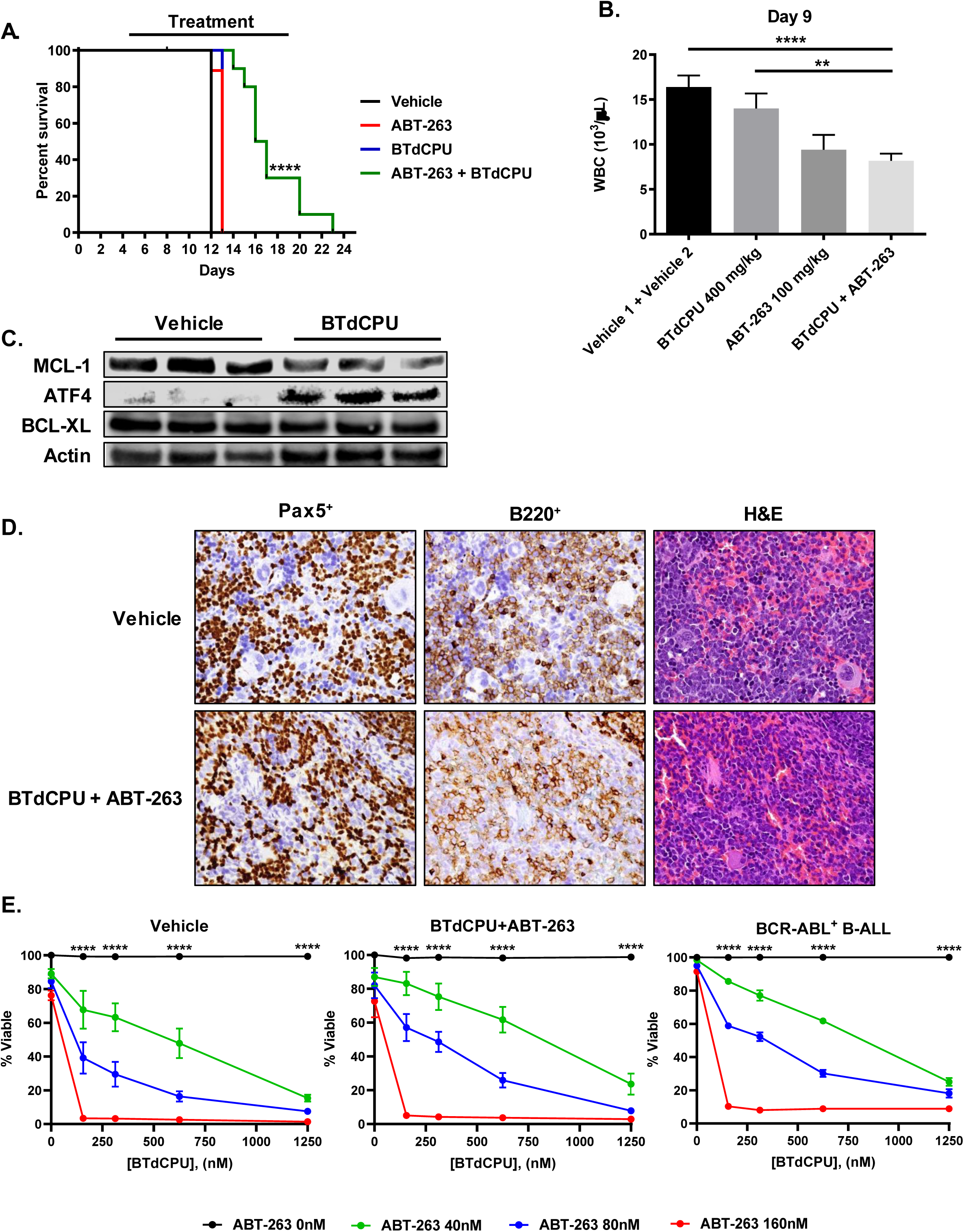
HRI activation synergizes with BH3-mimetics in vivo. Mouse BCR-ABL^+^ B-ALL cells were injected into unconditioned C57/BL6 recipients. After 5 days, the mice were divided to 4 treatment groups: vehicle, BTdCPU alone (400 mg/kg), ABT-263 alone (100 mg/kg), and combined BTdCPU+ABT-263. Mice were treated daily for 14 days. (**A**) Kaplan-Meier survival curve the cohorts of mice (n=9 at least for each treatment group). Log-rank test shows p<0.0001**** for combined treatment group. (**B**) Total white blood cell (WBC) count in peripheral blood of mice treated with the indicated groups on day 9 after leukemia injection. Each bar represents the average WBC count of the indicated group. Error bars indicate the standard error of mean at least 9 mice per group. Unpaired t-test indicates significance between vehicle and combined treatment group p<0.0001****. (**C**) Bone marrow from recipient mice treated with either vehicle or BTdCPU was harvested and immunoblotted for indicated proteins. (**D**) Immunohistochemistry from spleens of mice bearing BCR-ABL^+^ B-ALL treated with either vehicle or BTdCPU + ABT-263 at the time of sacrifice, stained as indicated. **(E)** Bone marrow harvested from mice treated with either vehicle or the combination of BTdCPU and ABT-263 was harvested and re-cultured with BTdCPU, ABT-263, or a combination of the two. After 24h, viability was measured by Annexin-V and propidium iodide staining. Two-way ANOVA with Bonferroni multiple comparison indicates significance P<0.0001**** between DHA or BTdCPU alone (0 nM ABT-263 or ABT-199) and 160 nM ABT-263 or ABT-199 at indicated doses of DHA or BTdCPU. The combination of BTdCPU+ABT-263 showed synergistic response on cells recovered from mice treated with vehicle (α=0.863, p=3.05e-10), BTdCPU+ABT-263 (α=2.58, p=2.36e-01) and in BCR-ABL^+^ B-ALL cells (α=0.697, p=1.39e-13).

Mechanistically, BTdCPU triggered the repression of MCL-1 in cultured human and mouse leukemia. We thus investigated whether BTdCPU could induce the repression of MCL-1 expression in mice by treating leukemia bearing recipient mice daily with BTdCPU. After 8 days of treatment the bone marrow was isolated, lysed, and assessed for protein expression by immunoblot analyses. The expression of MCL-1 in the bone marrow from leukemic mice treated with BTdCPU was significantly decreased when compared to that from mice treated with vehicle alone, whereas BCL-XL expression was unaffected by BTdCPU treatment (Fig. 6C, Sup. Fig. 6A). Furthermore, cells from mice treated with BTdCPU also exhibited elevated expression of ATF4 indicating the eIF2α pathway was activated (Fig. 6C).

Despite 14 days of treatment, the mice from the combination treatment group eventually succumbed to fatal leukemia as determined by immunohistochemical staining for expression of Pax5 and B220 on B-ALL cells (Fig. 6D). We therefore investigated if the leukemia cells in mice treated with the combination of BTdCPU and ABT-263 had acquired resistance to treatment. To complete this analysis, bone marrow from vehicle or combination treated mice were harvested, expanded, and then re-treated in culture in a synergy assay. Leukemia cells isolated from both mice treated with vehicle and the combination of BTdCPU and ABT-263 still showed a synergistic response to the treatment when treated *ex vivo*, indicating that the leukemia had not acquired resistance during the *in vivo* treatment (Fig. 6E and Sup. Fig. 6B). Taken together, these data indicate that activation of the HRI signal transduction pathway *in vivo* by BTdCPU can slow leukemia progression when combined with BH3-mimetic agents by repressing MCL-1 expression.

## Discussion

The observation that *MCL1* is frequently amplified in human cancers has driven significant efforts to develop potent and specific MCL-1 inhibitors (9). Indeed, several candidate MCL-1 inhibitors have shown on-target efficacy when tested in human hematological malignancies in culture and xenograft models (17–19, 45). Several of these agents are in early clinical trials. Despite some promising results in patients, in the fall of 2019 the FDA placed a clinical hold on the phase I trial of one MCL-1 inhibitor (AMG397) due to evidence of cardiac toxicity. In addition, mouse genetic evidence has shown that ablation of *Mcl1* in cardiomyocytes induces fatal cardiomyopathy (20, 21). These studies suggest that the direct inhibition of MCL-1 may have safety concerns in the clinic, especially when combined with standard chemotherapy that causes significant cardiac toxicity; highlighting the importance of discovering alternative mechanisms of targeting MCL-1.

Previous work from our laboratory identified that the widely used, anti-malarial drug dihydroartemisinin (DHA) triggers the repression of MCL-1 expression in mouse and human leukemic cells and synergizes with ABT-263 in culture and *in vivo* (22). Using both genetic and pharmacological approaches, here we provide mechanistic insight into how DHA induces the repression of MCL-1 translation. These efforts have revealed for the first time how a cellular heme-sensing pathway controls apoptotic sensitivity through the function of the HRI eIF2α kinase. In mouse and human leukemia, treatment with DHA triggers the HRI-dependent integrated stress response which results in eIF2α phosphorylation and represses cap-dependent translation. Since MCL-1 protein is labile and undergoes rapid basal proteasome-dependent turnover, inhibiting new protein translation rapidly results in the loss of MCL-1 expression (46). Here we demonstrate that loss of HRI in mouse and human leukemia cells blocks the DHA induced MCL-1 repression as well as the induction of downstream effectors ATF4 and CHOP. These data clearly implicate the HRI heme-sensing pathway as a key trigger of the integrated stress response resulting in the inhibition of MCL-1 translation.

MCL-1 is a known resistance factor for ABT-199 and ABT-263 BH3-mimetics, but DHA can repress MCL-1 expression in mouse and human leukemia, thus rendering the cells more sensitive to treatment with either ABT-199 or ABT-263. Importantly, in both mouse and human leukemia the loss of HRI significantly reduced the synergistic response to these BH3-mimetics when combined with DHA demonstrating the impact of this heme-sensing pathway on apoptotic sensitivity. These data led us to wonder if other pharmacological activators of HRI could similarly repress MCL-1 expression. To address this, we tested BTdCPU, a small molecule activator of HRI (43). Like DHA, BTdCPU, lead to MCL-1 repression and synergized with ABT-199 and ABT-263 BH3-mimetics in mouse and human leukemia. Consistent with *in vitro* data, we demonstrated that this pathway also repressed MCL-1 *in vivo* without any overt toxicity. This confirms the idea that activation of this heme-sensing pathway could be used to target MCL-1 and promote leukemic priming to BH3-mimetics.

The concept that perturbations of the heme synthesis pathway can affect apoptotic responses has precedent. In a metabolically focused CRISPR loss-of-function screen, acute myeloid leukemia (AML) cell lines were sensitized to ABT-199 induced killing when several heme synthesis genes were ablated (47). In this report, Wood and colleagues concluded that loss of heme synthesis exerts its apoptosis-sensitizing effects largely by disruption of electron transport chain function and loss of mitochondrial outer membrane integrity (47). However, precisely how disruptions in electron transport chain function contribute to permeabilization of the outer mitochondrial membrane was not clear from this study. Considering our observations, we postulate that defects in heme synthesis will, over time, lead to the activation of an HRI-dependent integrated stress response that would increase apoptotic priming by repressing MCL-1 expression. This loss of MCL-1 would promote cancer cell responses to BH3-mimetics such as ABT-199.

We reveal that the anti-malarial DHA requires heme synthesis to induce the HRI-dependent integrated stress response. In contrast, Wood and colleagues showed that blocking heme synthesis sensitized AML cell lines (47). However, we demonstrate that in a cell-free system DHA can physically disrupt the interaction between heme and HRI. Taken together, these data suggest that there may be a reaction between cellular heme and DHA itself that liberates HRI. Precisely how DHA disrupts the HRI and heme interaction, and why this process requires cellular heme is still unclear and will be the topic of further investigation. By better understanding the mechanisms of heme interactions and HRI activation one could imagine that it should be possible to develop more specific and potent drugs to target this pathway that could be used to treat a variety of types of malignancies.

Ph^+^ ALL has seen improvements in treatment; however, there is still a need for new treatment strategies as the intensive regimens have significant side-effects and are not effective in all patients (3, 4). Our work implicates an alternate therapeutic pathway that can be targeted in this subtype of leukemia. Perhaps even more importantly, we have shown that this pathway can also be targeted in Ph-like ALL, a harder to treat subtype known to have a poorer prognosis (5). We observed MCL-1 repression by both DHA and BTdCPU in mouse cell lines as well as human leukemia cells from both Ph^+^ and Ph-like ALL models. To continue to evaluate the potential of this pathway, we look forward to further validating these findings by combining DHA or BTdCPU with BH3-mimetics in NSG recipients bearing Ph^+^ or Ph-like primary patient leukemia.

We are interested in determining which other types of cancers utilize this HRI-dependent heme-sensing pathway for regulating apoptotic sensitivity. Since cancer cells frequently have elevated levels of heme and often exhibit up-regulated expression of many of the genes involved in the biosynthesis of heme (48), it is likely that our observations will be generalizable to cancer beyond B lineage acute leukemia. Indeed, inhibition of heme synthesis has even been reported to reduce tumor cell survival and proliferation in a variety of cancer types (30, 49, 50). The effects of inhibiting heme synthesis may be in part due to disruption of normal cellular processes including electron transport chain function, cataplerosis in the TCA cycle, p53 activity and stability, regulating trafficking of ADP and ATP, and in circadian rhythms (48). However, our novel findings have revealed mechanistically how activation of the cellular heme-sensing pathway can regulate apoptotic priming in leukemia.

It is too early to tell how much the development of potent and selective BH3-mimetic inhibitors of MCL-1 will continued to be hampered by toxic effects induced by MCL-1 inhibition. Despite these setbacks, MCL-1 remains an important therapeutic target for cancer therapy (9). Our data indicate that simply repressing MCL-1 expression by DHA or BTdCPU does not trigger any overt toxicities when used in mouse models (22). It is conceivable that partially repressing MCL-1 expression may cause a less severe insult to normal tissues than potent, pharmacological inhibition and thus, may avoid the associated toxicities. Therefore, we speculate that activation of this HRI-dependent heme-sensing pathway could be an effective and well-tolerated alternative approach to repress MCL-1 expression and render cancer cells more susceptible to approved BH3-mimetics.

## Authors’ Contributions

**Concept and design:** K.H. Smith, A. Budhraja, J.T. Opferman

**Development of methodology:** K.H. Smith, A. Budhraja, M.E. Turnis., J.P. Connelly, J. Lynch, J. Schuetz, S.M. Pruett-Miller, J.T. Opferman

**Acquisition of data (provided animals, etc.):** K.H. Smith, A. Budhraja, J. Lynch, J. Schuetz, S.M. Pruett-Miller, C.G. Mullighan, J.T. Opferman

**Analysis and interpretation of data:** K.H. Smith, A. Budhraja, M.E. Turnis, J.C. Panetta, J.P. Connelly, J. Lynch, J. Schuetz, S.M. Pruett-Miller, J.T. Opferman.

**Writing and review:** K.H. Smith, A. Budhraja, M.E. Turnis., J.T. Opferman

**Administrative, technical, or material support:** M.E. Turnis, K. Roberts

**Study Supervision:** K.H. Smith, A. Budhraja, J.T. Opferman

## Acknowledgments

We thank J.-J. Chen (MIT) for providing anti-HRI purified antibody and J. Slavish and Z. Rankovic (Department of Chemical Biology and Therapeutics, SJCRH) for synthesis of the small molecule, BTdCPU. We also thank S. Porter, S. Peters, S. Sakurada (Center for Advanced Genome Editing, SJCRH) for generation of mutant cell lines.

## Supplementary Tables

**Sup. Table 1:**
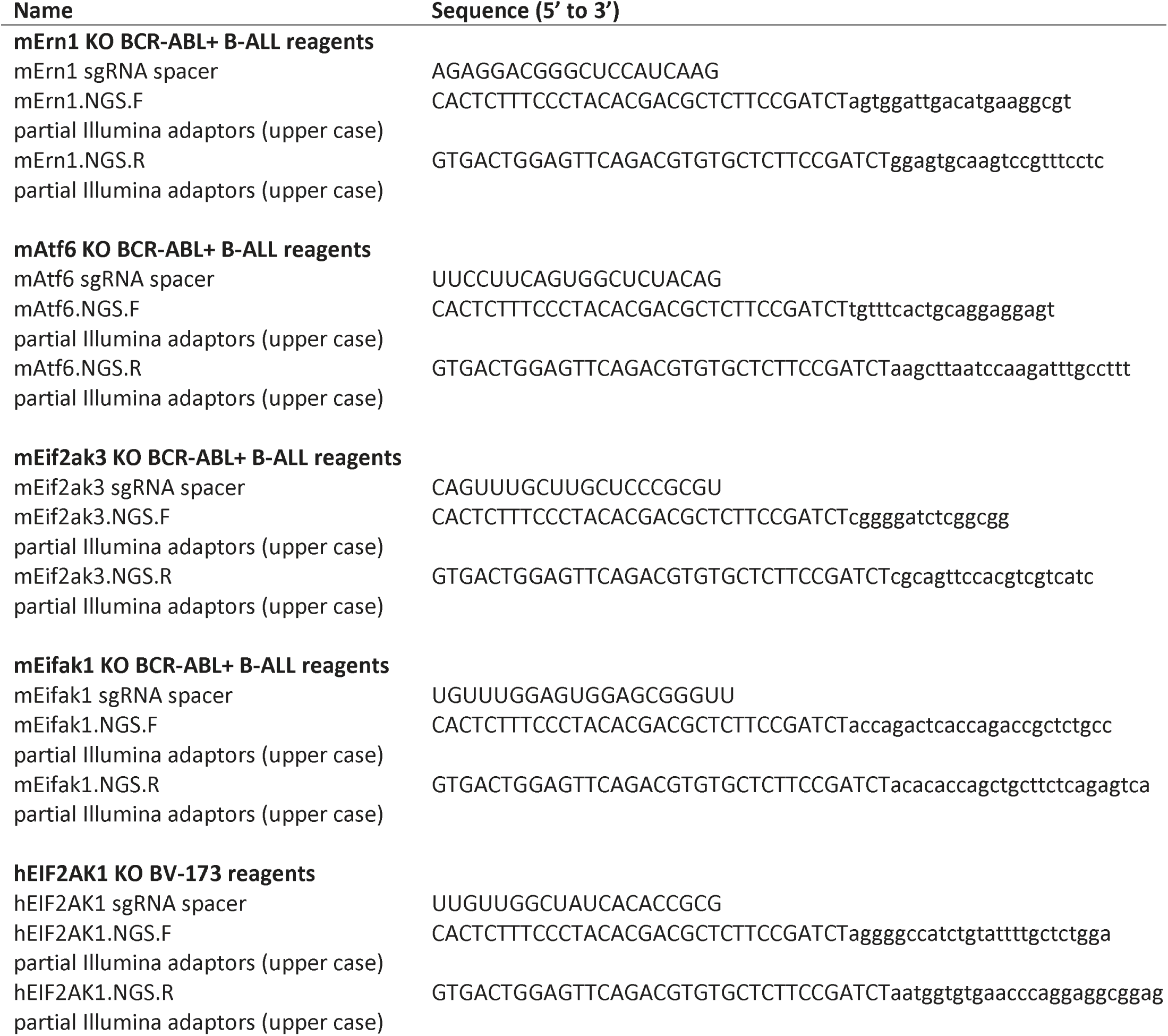
sgRNA sequences for knockout cell lines.

**Sup. Table 2:** Synergy analysis of combinatorial drug treatments, determined by response surface modeling. α-value: interaction term; drug combination is considered synergistic if the α-value is positive.

| Cell Type | Drug combination | α-value | p-value | Figure |
| --- | --- | --- | --- | --- |
| Tom1 | DHA + ABT-263 | 343 | 1.83E-28 | S4C |
| Tom1 | DHA + ABT-199 | 6.70E+03 | 8.28E-26 | S4C |
| Tom1 | BTdCPU + ABT-263 | 2.37 | 5.87E-07 | S4C |
| Tom1 | BTdCPU + ABT-199 | 337 | 2.41E-45 | S4C |
| OP1 | DHA + ABT-263 | 1.32E+05 | 1.49E-17 | S4D |
| OP1 | DHA + ABT-199 | 6.33E+03 | 3.72E-12 | S4D |
| OP1 | BTdCPU + ABT-263 | 4.84E+03 | 4.24E-17 | S4D |
| OP1 | BTdCPU + ABT-199 | 1.14E+03 | 1.74E-35 | S4D |
| SUP-B15 | DHA + ABT-263 | 565 | 2.02E-09 | S4E |
| SUP-B15 | DHA + ABT-199 | 404 | 1.68E-22 | S4E |
| SUP-B15 | BTdCPU + ABT-263 | 22.7 | 2.70E-08 | S4E |
| SUP-B15 | BTdCPU + ABT-199 | 5.81 | 4.47E-07 | S4E |
| Ba/F3 *RCSD1-ABL1* | DHA + ABT-263 | 484 | 2.45E-45 | S4H |
| Ba/F3 *RCSD1-ABL1* | DHA + ABT-199 | 472 | 1.78E-38 | S4H |
| Ba/F3 *RCSD1-ABL1* | BTdCPU + ABT-263 | 105 | 3.85E-60 | S4H |
| Ba/F3 *RCSD1-ABL1* | BTdCPU + ABT-199 | 126 | 5.70E-54 | S4H |
| Ba/F3 *RCSD1-ABL2* | DHA + ABT-263 | 1.15E+03 | 2.02E-47 | S4I |
| Ba/F3 *RCSD1-ABL2* | DHA + ABT-199 | 951 | 4.47E-46 | S4I |
| Ba/F3 *RCSD1-ABL2* | BTdCPU + ABT-263 | 153 | 2.45E-74 | S4I |
| Ba/F3 *RCSD1-ABL2* | BTdCPU + ABT-199 | 179 | 2.71E-57 | S4I |

## Supplemental Figure Legends

**Figure S1.**
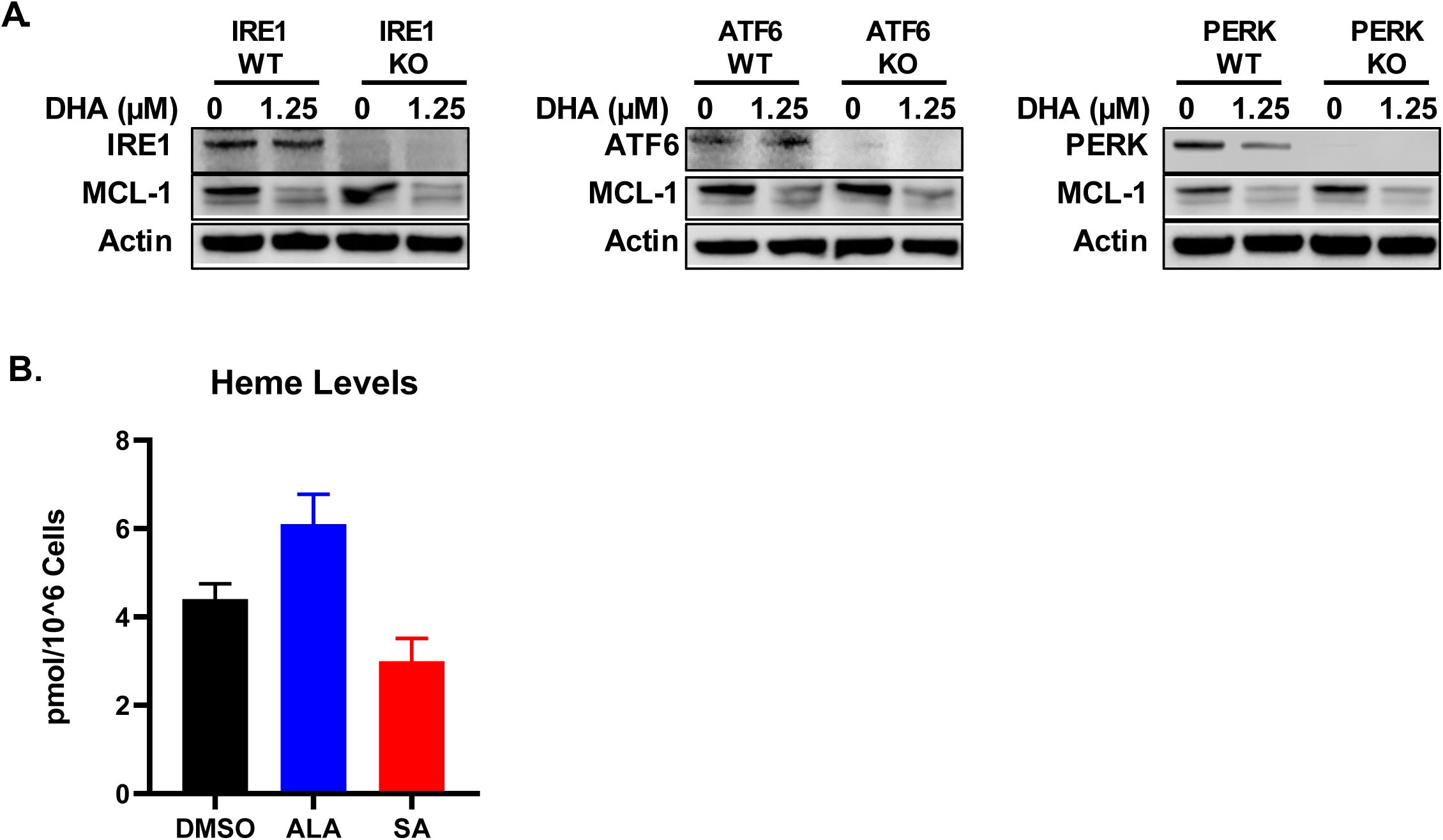
Stress pathways in cellular response to DHA. **(A)** Wild-type (WT) or BCR-ABL^+^ B-ALL cells deficient (KO) for *ERN1* (encodes IRE1), *ATF6*, or *Eif2ak3 (*encodes PERK) were treated with 1.25µM DHA for 9h and MCL-1 expression was determined by immunoblotting with indicated antibodies. **(B)** Heme levels were measured in BCR-ABL^+^ B-ALL cells treated with DMSO, 62.5µM (Aminolevulinic acid) ALA, or 62.5µM (succinylacetone) SA for 24h.

**Figure S2.**
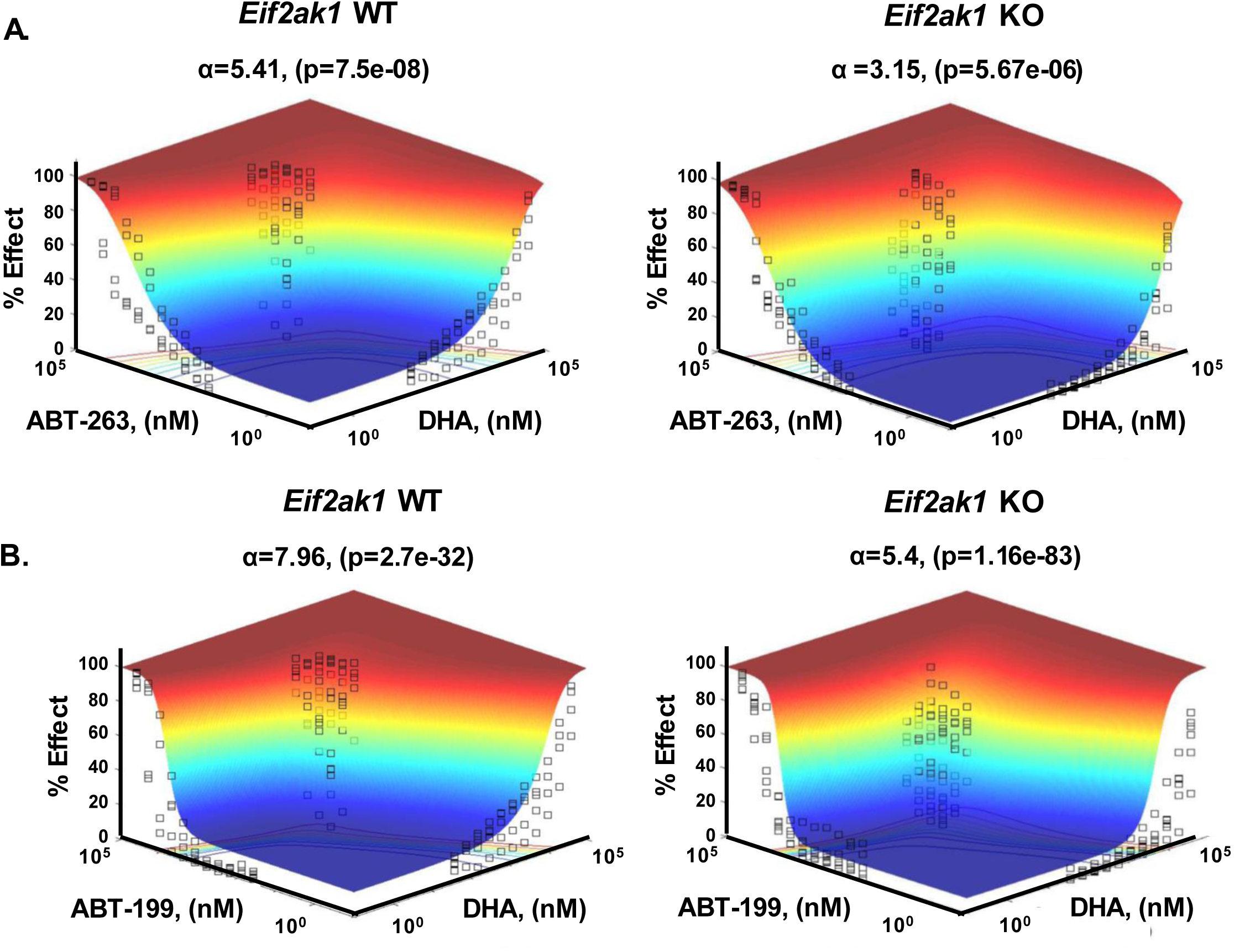
Synergy analysis of DHA and BH3-mimetics in BCR-ABL^+^ B-ALL cells. Response surface modeling was used to assess synergy in wild-type (WT) or *Eif2ak1*-KO (HRI-deficient) BCR-ABL^+^ B-ALL cells. **(A)** The combination of DHA and ABT-263 showed less synergistic response in *Eif2ak1*-KO (α=3.15, p=5.67e-06) as compared to wild-type (α=5.41, p=7.5e-08) BCR-ABL B-ALL cells. Difference in α-values between wild-type and *Eif2ak1*-KO were statistically significant (p<10^-5^). **(B)** The combination of DHA and ABT-199 showed less synergistic response in *Eif2ak1*-KO (α=5.4, p=1.16e-83) as compared to wild-type (α=7.96, p=2.7e-32) BCR-ABL B-ALL cells. Difference in α-values between wild-type and *Eif2ak1*-KO were statistically significant (p<0.057).

**Figure S3.**
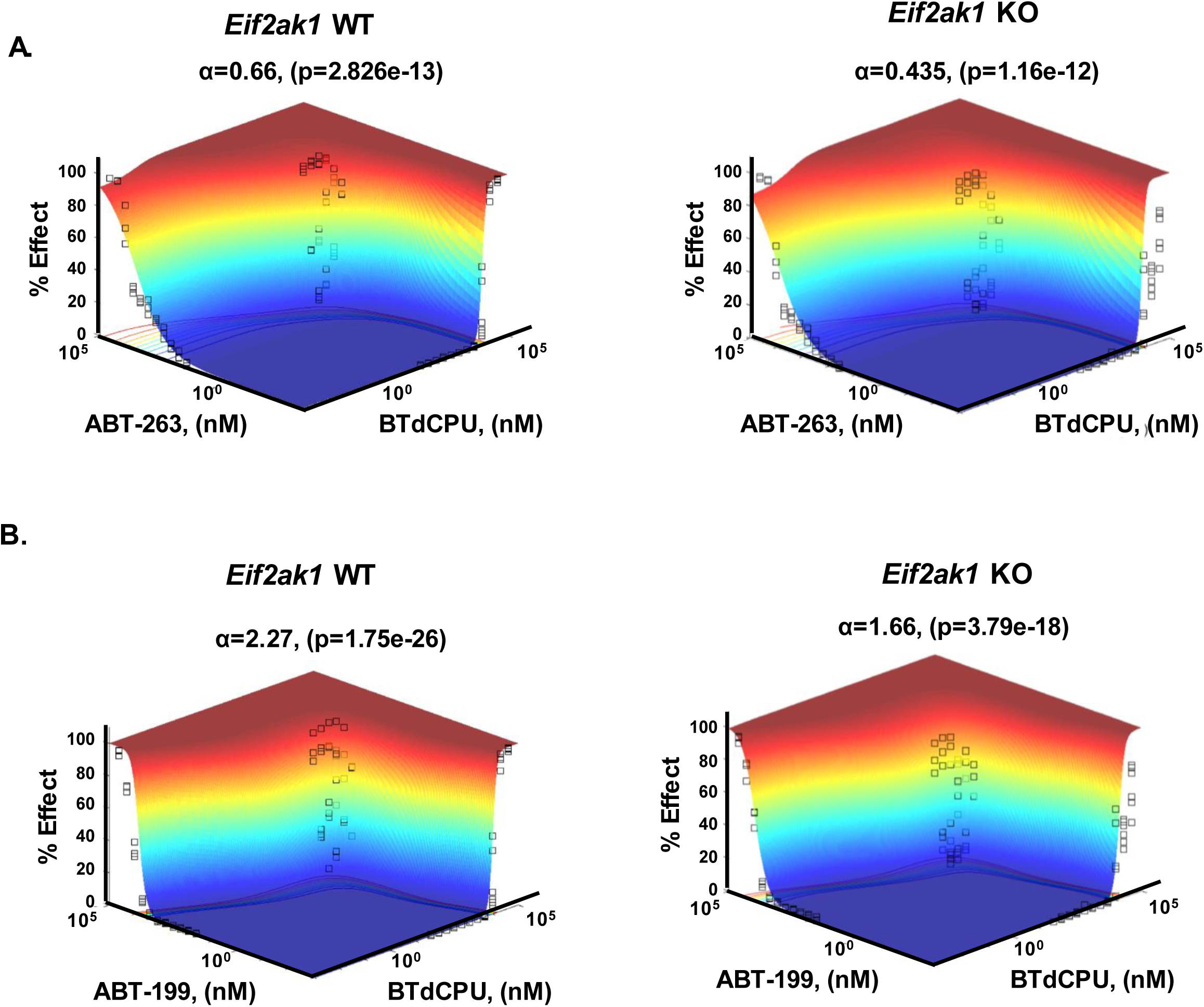
Synergy analysis of BTdCPU and BH3-mimetics in BCR-ABL^+^ B-ALL cells. Response surface modeling was used to assess synergy in wild-type (WT) or *Eif2ak1*-KO (HRI-deficient) BCR-ABL^+^ B-ALL cells. **(A)** The combination of BTdCPU and ABT-263 showed less synergistic response in *Eif2ak1*-KO (α=0.435, p=1.16e-12) as compared to wild-type (α=0.66, p=2.826e-13) BCR-ABL^+^ B-ALL cells. Difference in α-values between wild-type and *Eif2ak1*-KO were statistically significant (p<0.05). **(B)** The combination of BTdCPU and ABT-199 showed less synergistic response in *Eif2ak1*-KO (α=1.66, p=3.79e-18) as compared to wild-type (α=2.27, p=1.75e-26) BCR-ABL B-ALL cells. Difference in α-values between wild-type and *Eif2ak1*-KO were statistically significant (p<0.05).

**Figure S4.**
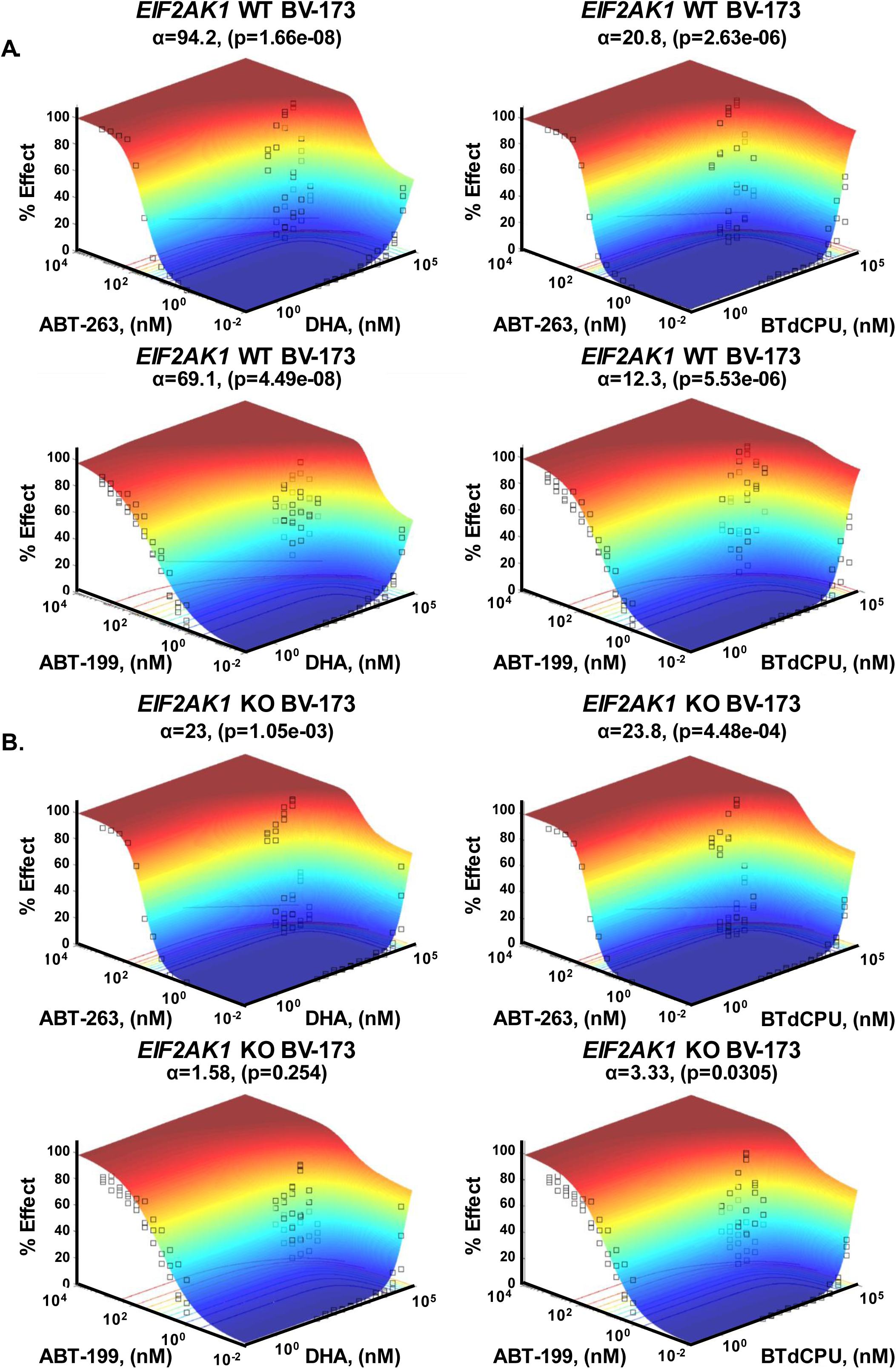
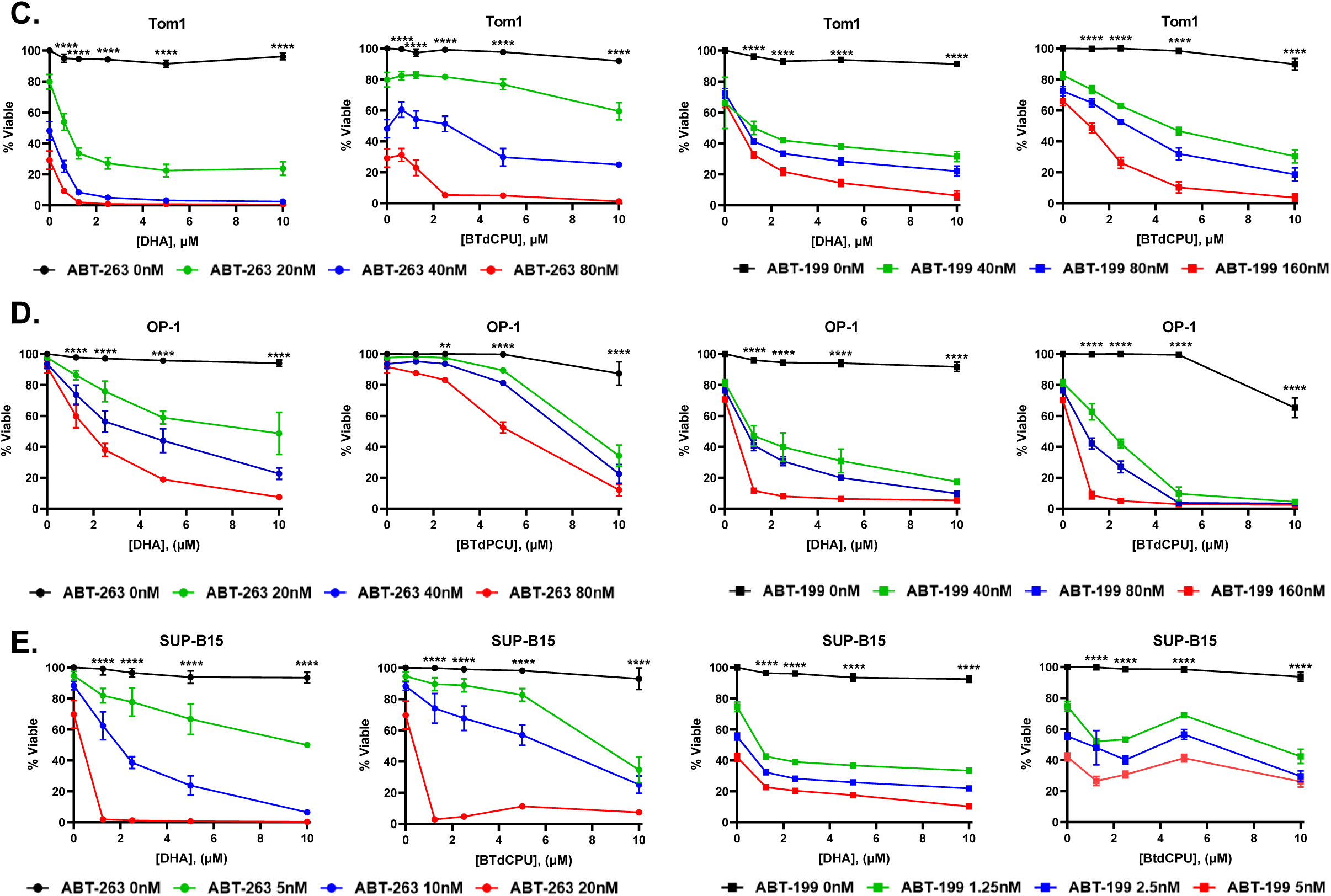
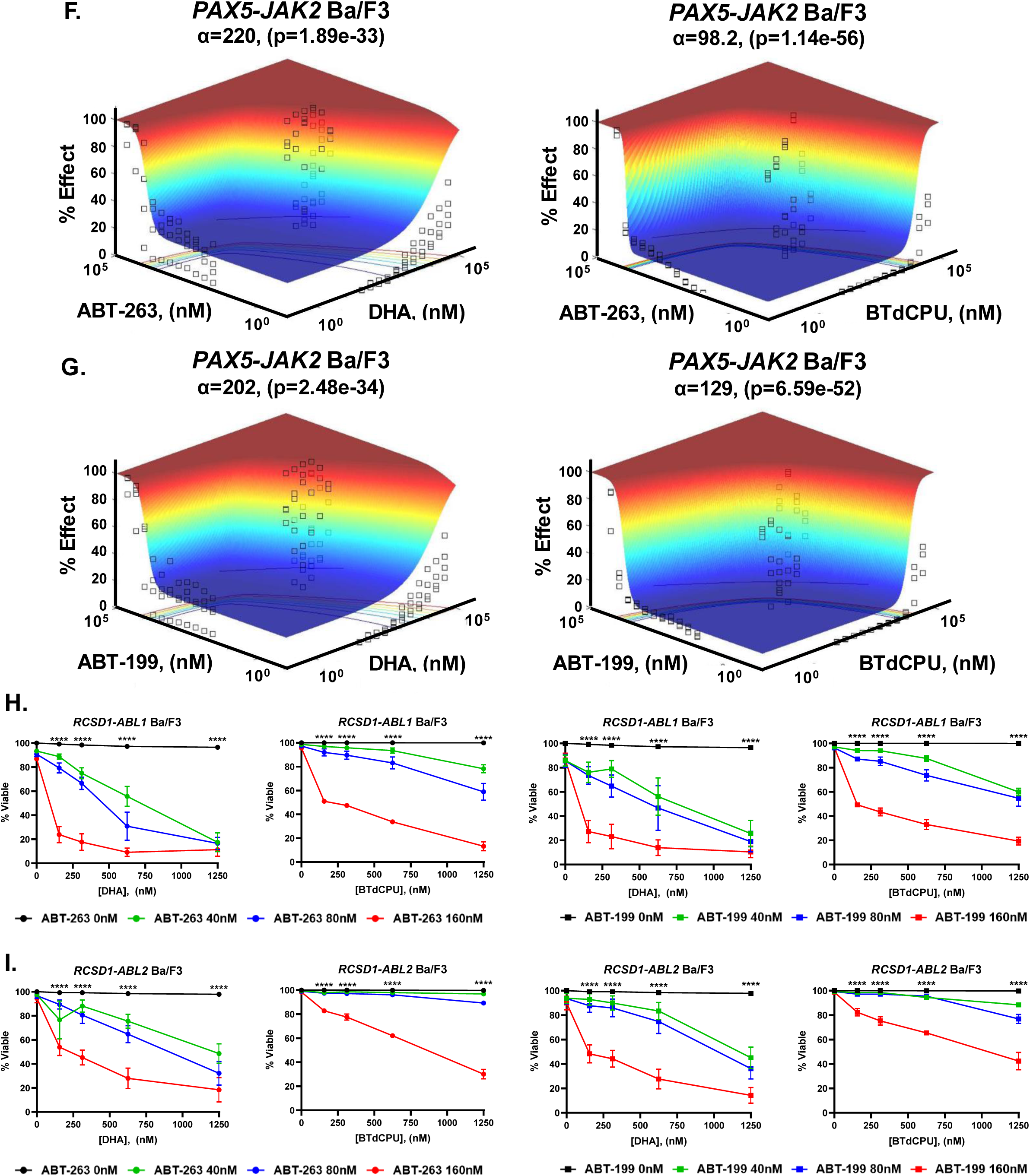
HRI activation synergizes with BH3-mimetics in Ph^+^ and Ph-like ALL cell lines. Response surface modeling was used to assess synergy in **(A)** wild-type (WT) or **(B)** *EIF2AK1*-KO (HRI-deficient) BV173 cells treated with DHA or BTdCPU combined with ABT-263 or ABT-199. The combinations of DHA with ABT-199 or ABT-263 and BTdCPU with ABT-199 were more synergistic in WT cells compared to *EIF2AK1-*KO cells (the differences in alpha values were significant (p<10^-3^)). **(C-E)** Human Ph^+^ ALL cell lines **(C)** Tom1, **(D)** OP-1, and **(E)** SUP-B15 were treated with the indicated concentrations of ABT-263 or ABT-199 combined with DHA or BTdCPU for 24h and viability was measure by Annexin-V and propidium iodide staining. Two-way ANOVA with Bonferroni multiple comparison indicates significance P<0.0001**** between DHA or BTdCPU alone (0 nM ABT-263 or ABT-199) and 80 nM ABT-263 or 160 nM ABT-199 **(for C and D)** and 20 nM ABT-263 or 5 nM ABT-199 **(for E)** at indicated doses of DHA or BTdCPU. **(F and G)** Response surface modeling was used to assess synergy of these drug combinations, indicated by an α value. Response surface modeling was used to assess synergy in *PAX5-JAK2* Ba/F3 cells treated with DHA or BTdCPU combined with **(F)** ABT-263 or **(G)** ABT-199. **(H and I)** Ba/F3 cells expressing the **(H)** *RCSD1-ABL1* or **(I)** *RCSD1-ABL2* fusions were treated with 0 nM, 40 nM, 80 nM, or 160 nM of ABT-263 or ABT-199 combined with DHA or BTdCPU for 24h and viability was measured by Annexin-V and propidium iodide staining. Response surface modeling was used to assess synergy of these drug combinations, indicated by an α value. Two-way ANOVA with Bonferroni multiple comparison indicates significance P<0.0001**** between DHA or BTdCPU alone (0 nM ABT-263 or ABT-199) and 160 nM ABT-263 or ABT-199 at indicated doses of DHA or BTdCPU. See Supplementary Table 2 for alpha values.

**Figure S5.**
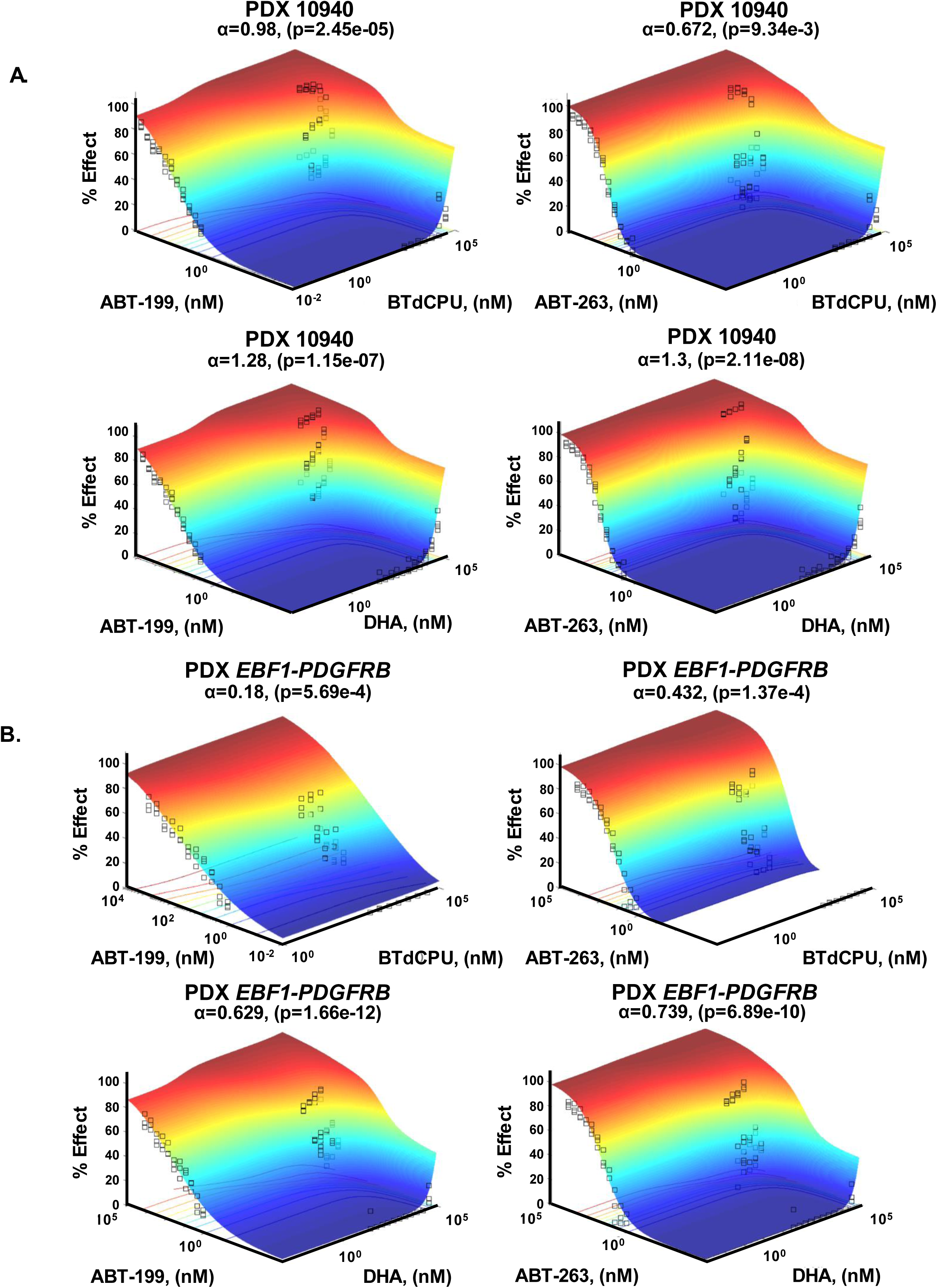
Synergy analysis of DHA or BTdPCU combined with BH3-mimetics in PDX cells. **(A and B)** Response surface modeling was used to assess synergy in **(A)** Ph^+^ 10940 or **(B)** *EBF1-PDGFRB* Ph-like PDX cells treated with DHA or BTdCPU combined with ABT-263 or ABT-199. See Supplementary Table 2 for alpha values.

**Figure S6.**
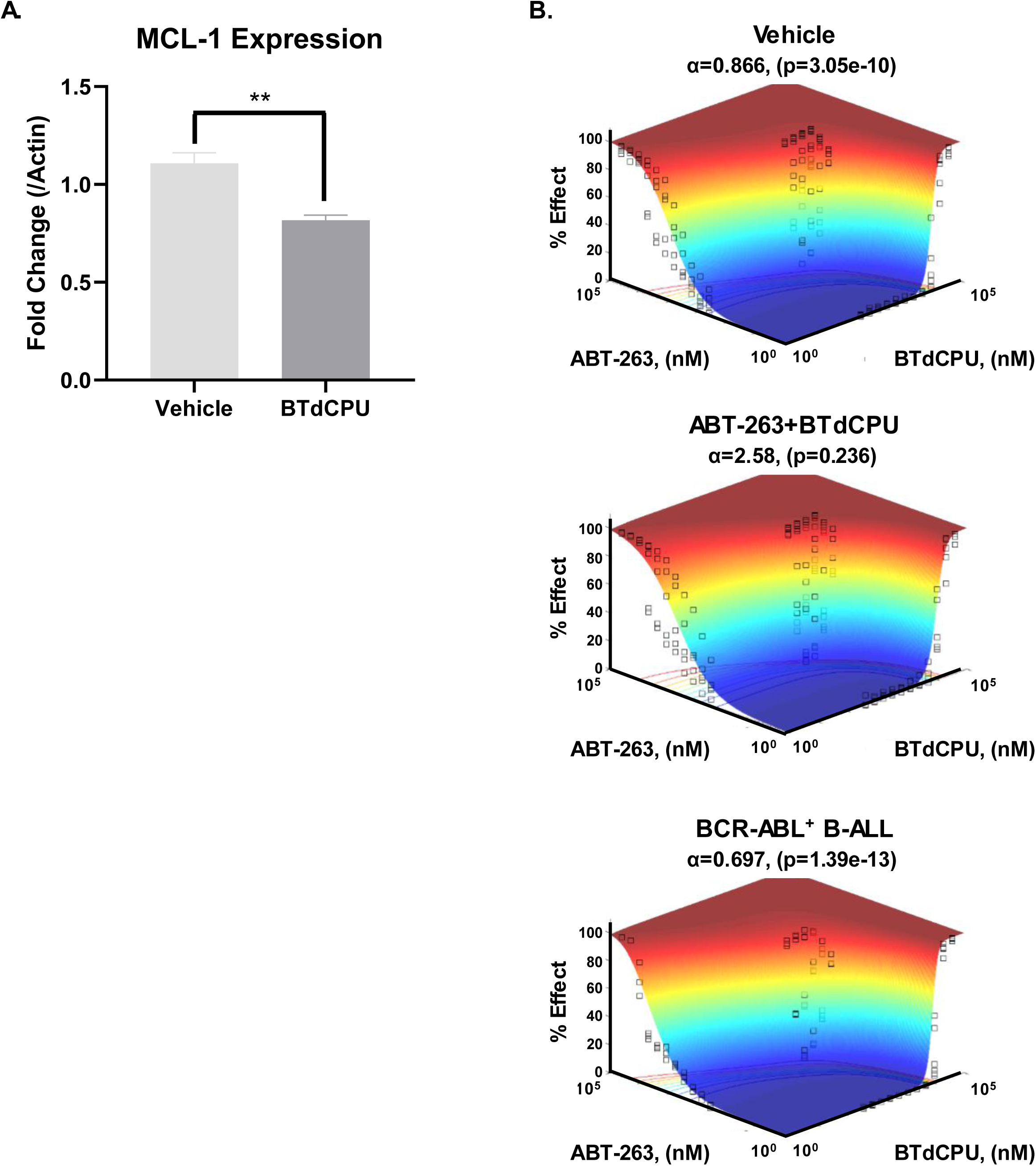
Densitometry and synergy analysis of HRI activation *ex vivo*. **(A)** Densitometry of MCL-1 expression in bone marrow cells harvested from BTdCPU treated mice (related to Fig. 6C). Data are the average of 3 separate animals and error bars are SEM. Unpaired t-test indicates significance between vehicle and BTdCPU p<0.01**. **(B)** Response surface modeling was used to assess synergy in cells isolated from vehicle or combination treated mice when treated with BTdCPU and ABT-263 *ex vivo*.

